# Gliotransmission of D-serine promotes thirst-directed behaviors in *Drosophila*

**DOI:** 10.1101/2022.03.07.483255

**Authors:** Annie Park, Vincent Croset, Nils Otto, Devika Agarwal, Christoph D. Treiber, Eleanora Meschi, David Sims, Scott Waddell

## Abstract

Thirst emerges from a range of cellular changes that ultimately motivate an animal to consume water. Although thirst-responsive neuronal signals have been reported, the full complement of brain responses is unclear. Here we identify molecular and cellular adaptations in the brain using single-cell sequencing of water deprived *Drosophila*. Water deficiency primarily altered the glial transcriptome. Screening the regulated genes revealed astrocytic expression of the *astray*-encoded phosphoserine phosphatase to bi-directionally regulate water consumption. Astray synthesizes the gliotransmitter D-serine and vesicular release from astrocytes is required for drinking. Moreover, dietary D-serine rescues *aay*-dependent drinking deficits while facilitating water consumption and expression of water-seeking memory. D-serine action requires binding to neuronal NMDA-type glutamate receptors. Fly astrocytes contribute processes to tripartite synapses and the proportion of astrocytes that are themselves activated by glutamate increases with water deprivation. We propose that thirst elevates astrocytic D-serine release, which awakens quiescent glutamatergic circuits to enhance water procurement.

## Introduction

The sensation of thirst is predominantly a manifestation of our body’s response to being water deprived. Water deficit reduces the volume and increases the osmolarity of an animal’s blood. These changes induce multiple adaptations throughout the body, such as elevated activity of osmosensory neurons in the brain, alteration of blood pressure and heart rate, and increased retention of water.

Osmosensory neurons in the subfornical organ (SFO) and organum vasculosum (OVLT) of the lamina terminalis (LT) of the mammalian forebrain respond directly and indirectly to changes in plasma osmolality and blood volume/pressure, via hormones such as Angiotensin II (Bourque, 2008; Fitzsimons, 1998; Pool et al., 2020). These neurons then directly or indirectly release compensatory neuropeptide/hormone signals like Vasopressin/anti-diuretic hormone which promotes bodily responses to reduce water loss and restore blood pressure, and behavioral responses to restore fluid balance (Augustine et al., 2020; Johnson and Thunhorst, 1997; Rolls, 1971; Zimmerman et al., 2017). Artificial engagement of LT neurons can induce water consummatory behaviors (Abbott et al., 2016; Andersson and McCann, 1955; Augustine et al., 2018; Betley et al., 2015; Chen et al., 2016; Gizowski et al., 2016; Matsuda et al., 2017; Oka et al., 2015; Zimmerman et al., 2016, 2019). However, the full range of nervous system mechanisms that accompany the water-deprived state, and how they modulate behavioral regimes and actions to satisfy thirst, is currently unknown.

In *Drosophila* Ion Transport Peptide (ITP) is the likely antidiuretic functional analog of the mammalian vasopressin and renin-angiotensin systems (Gáliková et al., 2018). ITP is produced by neurosecretory neurons in the brain and abdominal ganglion. ITP expression increases with water-deprivation and it increases water consumption behavior, reduces water excretion in the malpighian tubules and increases reabsorption in the fly hind gut. ITP induction also represses feeding. Although neuronal circuits regulated by ITP remain to be identified, other thirst-regulated circuits that are required for water seeking and consumption/homeostasis have been reported. Interoceptive sensory neurons (ISNs) in the subesophageal zone (SEZ) directly sense high osmolality (via the cation channel Nanchung) and their inhibition promotes drinking (Jourjine et al., 2016). Interestingly, ISN activation (via Adipokinetic Hormine, AKH) suppresses drinking and instead promotes feeding. Two other classes of SEZ neurons, the Janu neurons, regulate water seeking behavior up a humidity gradient, but not total water consumption (Landayan et al., 2021). The Janu neurons are either GABAergic or Allatostatin (AstA)-releasing. Their joint activation is rewarding and the AstA group simultaneously inhibits feeding while promoting water seeking. Lastly, different types of mushroom body innervating dopaminergic neurons have been implicated in water-seeking, water reward learning (Lin et al., 2014b), and thirst-dependent control of the expression of water-seeking memories (Senapati et al., 2019). Some of these dopaminergic neurons are regulated by the leucokinin neuropeptide which is released from neurons that are activated by elevated osmolality (Senapati et al., 2019). More complex interaction between neuromodulators allows the dopaminergic neurons to selectively promote state appropriate expression of thirst- or hunger-dependent memories.

Brain responses to water-deprivation need not be exclusively neuronal. A blood-brain barrier (BBB) shields most neurons from the circulatory environment. Notably, the osmosensory neurons of the mammalian SFO and OV have processes outside the BBB where they can directly sample the circulatory status (Persidsky et al., 2006). The mammalian BBB is formed by microvascular endothelial cells, pericytes and astrocytes. Since astrocytes also innervate the neuropil, where their processes contribute to tripartite synapses, they are well positioned to sense and/or transmit metabolites and signals representing nutrient status to neurons (Fischer et al., 1997; Gourine and Kasparov, 2011; Limmer et al., 2014; Ronaldson and Davis, 2020; Simard and Nedergaard, 2004; Teschemacher et al., 2015; de Tredern et al., 2021). In *Drosophila*, perineurial and subperineurial glia form the BBB whereas astrocytes tile the entire brain and permeate the neuropil, where they associate with neuronal processes (Limmer et al., 2014; Stork et al., 2014). Astrocytes in the fly therefore also have potential to influence neuronal activity in response to nutritional state (Volkenhoff et al., 2015). However, evidence for such a role of fly astrocytes is currently lacking.

The *Drosophila* brain provides an unparalleled opportunity to investigate the neural mechanisms of thirst with cellular and molecular resolution. Recent advances in single-cell transcriptomics have enabled the generation of transcriptional profiles from most of the major cell types in the fly brain (Croset et al., 2018; Davie et al., 2018; Janssens et al., 2022; Konstantinides et al., 2018; Özel et al., 2021). Here, we applied this technology and knowledge to identify brain-wide and cell-type restricted changes in gene expression triggered by water-deprivation. Surprisingly, the majority of transcriptional changes occurred within glia, rather than neurons. Functional analyses of thirst responsive genes identified *astray* (*aay*) expression to be required in astrocytes for regulated water consumption. The *aay*-encoded phosphoserine phosphatase can produce the gliotransmitter D-serine. Astrocyte-specific suppression of *aay* reduced drinking in thirsty flies, while D-serine feeding restored these defects and facilitated water consumption in wild-type flies. Mutations modifying the D-serine binding site of fly NMDARs predictably altered water consumption. Connectomics revealed astrocytes to contribute processes to tripartite glutamatergic synapses and water deprivation increased the number of astrocytes that were responsive to glutamate. Our results support a model in which thirst increases astrocytic release of D-serine to recruit glutamatergic synaptic connections within neuronal circuits that promote water seeking and consumption. These findings provide a new molecular and cellular framework to understand how thirst alters brain physiology and behavior.

## Results

### Single-cell transcriptomics in thirsty *Drosophila*

*Drosophila* prefer dry environments when water-sated but seek humidity when water-deprived, behaviors that serve fluid homeostasis (Lin et al., 2014b; Liu et al., 2007; Sayeed and Benzer, 1996). This behavioral switch is readily quantified by giving flies the choice in a T-maze between a humid and a dry chamber (Figure 1A). Whereas water-sated flies preferred the dry chamber those dehydrated for 6h or 12h showed an increasing preference for the humid chamber (Figure 1B). Flies water-deprived for 12h then permitted to drink water reverted to preference for the dry chamber, demonstrating that water attraction is rapidly reversed upon drinking (Figure 1A and 1B).

**Figure 1.**
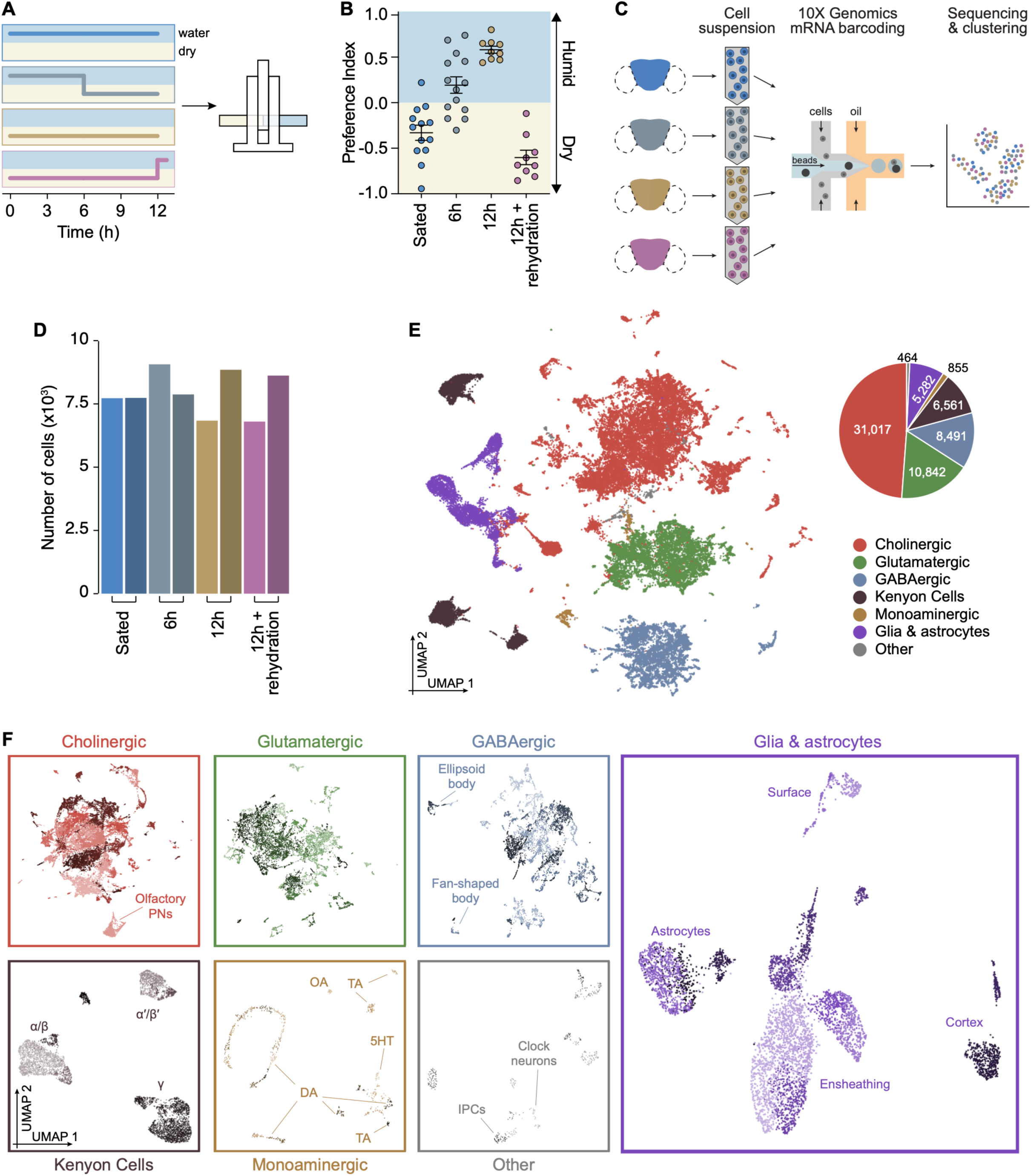
Single-cell transcriptomics of thirsty *Drosophila*. (**A**) Schematic protocols for humidity preference assay. Flies were kept in vials with or without water for the indicated times, then given a T-maze choice between a humid and a dry chamber. (**B**) Increasing dehydration converts humidity avoidance behavior into attraction. Attraction returns to avoidance with thirst quenching. (**C**) Schematic of process of single-cell transcriptomics analyses comparing flies from the four conditions in (**B**). Two independent samples were processed for each condition. (**D**) Total number of cells obtained from each sample, after filtration of low-quality barcodes and doublets. (**E**) Left: UMAP plot from first clustering step, and identification of seven main cell classes. Right: pie chart showing number of cells obtained from each of these classes. (**F**) UMAP plots showing sub-clustering of each of the seven cell classes shown in (**E**). Known cell types within each class are labelled. Kenyon Cell labels represent known subtypes that innervate corresponding mushroom body lobes. Withing the monoaminergic cells labels are OA: octopaminergic; TA: tyraminergic; 5HT: serotonergic; DA: dopaminergic. Within Other, IPC: insulin-producing cells. See also **Figure S1**.

To investigate cellular correlates of thirst we used the 10X Genomics Chromium system to generate single-cell transcriptomic atlases of brains extracted from water-sated, 6h dehydrated, 12h dehydrated, and rehydrated flies (Figure 1C). After filtering out low-quality barcodes and putative cell doublets (Figure S1A), we retrieved a total of 63,512 cells. Importantly, each dehydration condition and experimental sample contributed a similar number of cells to the total collection (Figure 1D). We used SCTransform and the canonical correlation anchor-based method in Seurat v3 (Hafemeister and Satija, 2019; Stuart et al., 2019) to normalize and integrate data from each sample to help identify shared cell populations. An initial unsupervised clustering step was performed to partition cells into seven main classes, which we annotated based on the expression of known marker genes (Figure S1B). These classes include cholinergic, glutamatergic, GABAergic, mushroom body intrinsic neurons or Kenyon Cells, monoaminergic neurons, glia, as well as “other” cells that did not fit into these previous categories (Figure 1E) (Barnstedt et al., 2016; Croset et al., 2018). Cells in each class were next independently subdivided into a total of 184 clusters each containing between 14 and 5083 cells (Figure 1F). Monoaminergic clusters were considerably smaller than others (median cells/cluster: 32), reflecting their discrete transmitters (DA, OA, TA, 5HT), the high functional specialization amongst neurons of each type, and their relative scarcity in the brain. Conversely, Kenyon Cell and other cholinergic neuron clusters were the largest (median cells/cluster: 354.5 and 293.5, respectively; Figure S1C). Here again, for each of these 184 clusters, we identified specific marker genes (Data S1, Figure S1F) which we used to annotate known cell types (Figure 1F; see Methods for detail). We noted that each experimental sample was evenly represented across cell clusters (Figure S1D and S1E), indicating that dehydration does not change the cellular composition of the mature adult fly brain, in contrast to a report from developing larvae where starvation reduced the number of undifferentiated *headcase* expressing neurons (Brunet Avalos et al., 2019).

### The transcriptional signature of thirst in the fly brain

A characteristic of single-cell RNA-seq (scRNA-seq) data is that expression of certain genes is stochastically undetectable in a fraction of the cells in which they are normally expressed. This artefact called ‘dropout’ can result from low mRNA levels in single cells and inefficient mRNA capture during library preparation, and it produces a larger than expected number of zero read counts (Hwang et al., 2018; Kharchenko et al., 2014). Therefore, classic differential expression methods designed for negative binomial bulk transcriptome data often underperform when analyzing weakly expressed genes in scRNA-seq data (Mou et al., 2020; Soneson and Robinson, 2018). To correct for potential bias caused by zero read count inflation we downweighed excess zeros using ZINB-WaVE (Risso et al., 2018; Van den Berge et al., 2018), which enables a more accurate estimation of data dispersion and thereby improves detection of differentially expressed genes. Using ZINB-WaVE with both edgeR and DESeq2, two established tools for quantifying differential expression (Love et al., 2014; Robinson et al., 2010), we first calculated gene expression differences between water-sated and 12h-dehydrated conditions, across cell clusters (Figure 2A).

**Figure 2.**
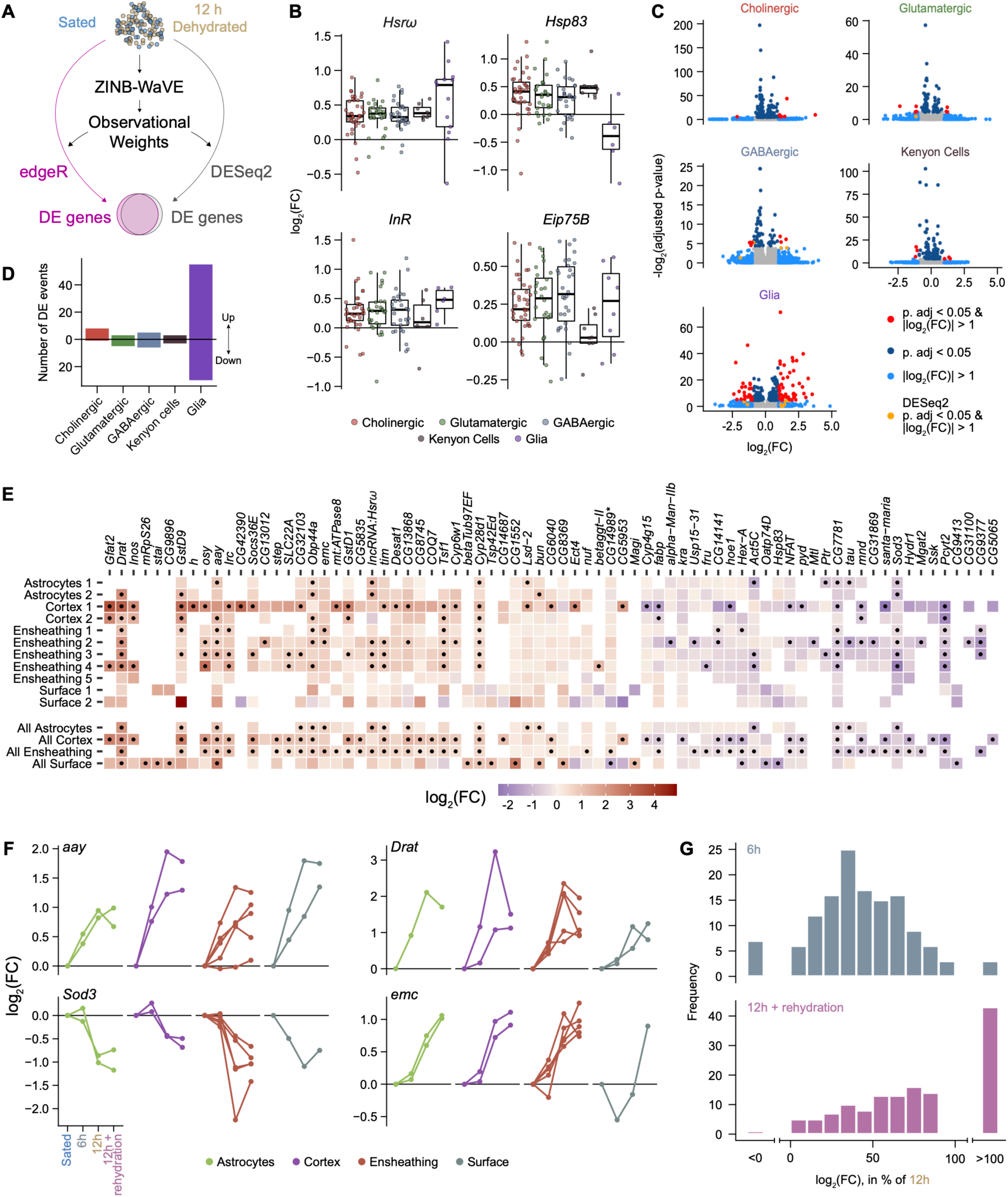
The transcriptional signature of thirst in the fly brain. (**A**) Schematic of differential expression analysis. Observational weights calculated with ZINB-WaVE were used in edgeR and DESeq2 to correct for zero-inflation. (**B**) Boxplots showing differential expression of the four genes most broadly regulated in each cluster after 12 h dehydration, grouped by main cell class. (**C**) Volcano plots representing statistical significance against fold change for all genes tested in cholinergic neurons, glutamatergic neurons, GABAergic neurons, Kenyon Cells and glia, calculated with edgeR. Each plot represents pooled data from all clusters in each cell class. Genes with adjusted p-value < 0.05 and |log2(FC)| > 1 in DESeq2 but not edgeR are labelled orange. (**D**) Number of differential expression events identified in the five cell classes shown in (**C**). Values above the bar represent genes up-regulated in thirsty flies, and below the bar represents down-regulated genes. (**E**) Heatmap showing fold change for each of the most regulated genes in glia, for each cluster (top) and for clusters grouped by glia type (bottom), calculated with edgeR. Black dots: adjusted p-value < 0.05. Empty tiles: transcript levels below detection threshold. *: *CG14989* is the only gene significantly up-and down-regulated in different clusters. (**F**) Changes in expression for four differentially expressed genes across glia clusters, and through all four hydration conditions. In most cases, expression gradually increases or decreases during dehydration. After 12 h re-hydration, mRNA levels tended to remain similar to levels measured in dehydrated flies. (**G**) Comparisons of log2(FC) values (vs. sated controls) after 6 h dehydration (top) or after rehydration (bottom), as a percentage of log2(FC) values after 12 h dehydration. Percentages were calculated for each of the significant gene-cluster pairs shown in (E) (black dots) and represented in histograms in 10% increments. Changes in expression are lower (<100%) after 6 h than after 12 h dehydration (top) for most genes. However, expression of many genes remains high (>100%) after thirsty flies are allowed to drink (bottom). See also **Figure S2**.

We expected that general osmotic stress resulting from dehydration might trigger a brain-wide transcriptional response. Consistent with this prediction we identified four genes whose expression increased broadly across most clusters, with an average log_2_(FC) across clusters above 0.25 (Figure 2B). Two of these, the chaperone encoding *Hsp83* and the long non-coding RNA *Hsrw*, have been previously implicated in the cellular stress response and shown to interact with each other to regulate transcription (Lakhotia, 2011; Qin et al., 2005; Ray et al., 2019). Global upregulation of the insulin receptor (*InR*) may reflect the role of insulin signaling in balancing thirst and hunger (Jourjine et al., 2016; Liu et al., 2015), whereas the ecdysone-response gene *Eip75B* was shown to protect circadian rhythms in stressful conditions (Kumar et al., 2014).

Cell-types that exhibit robust state-dependent changes in gene expression potentially play a role in the fly’s response to thirst, and the regulated genes may represent intra or intercellular molecular mechanisms that mediate physiological and behavioral responses. We found 84 genes that exhibited strong differential expression (|log_2_(FC)|>1, adjusted p-value <0.05) in at least one cell cluster, with either edgeR or DESeq2, which identified slightly different sets of genes. (Figure S2A). Most differential expression events (85/119) occurred in glia, with the rest in clusters from cholinergic, glutamatergic, GABAergic and Kenyon cell classes (Figures 2C and 2D). No individual cluster within the monoaminergic and “other” classes contained enough cells to enable differential analysis with sufficient statistical power. Differential expression was not apparent when clusters from these two classes were analyzed as a group. Within the glial clusters, 6 genes passed our differential expression criteria in astrocytes, 26 in ensheathing glia, and 33 in cortex glia. No differences were apparent in surface glia. However, pooling glial clusters by type provided enough statistical power to reveal 15 genes in surface glia and 2 additional genes in cortex glia. (Figure 2E). With the exception of *CG14989* which was simultaneously upregulated in ensheathing glia and downregulated in surface glia, all other differentially expressed genes changed in a similar direction in multiple glial cell types. Gene Ontology analysis indicated that the differentially expressed genes contribute to pathways related to metabolism, the response to stress, or behavior (Figure S2D). Together these results show that the highest magnitude thirst-dependent changes in gene expression occur in glial cell-types, whereas neuronal transcriptomes remain comparatively stable.

We next tested whether the expression of thirst-responsive genes changed with dehydration time. For several, including *aay* (*astray*), *Drat* (*Death resistor Adh domain containing target*) and *emc* (*extra macrochaetae*), expression steadily increased as dehydration extended from 0 to 6 to 12 h (Figure 2F), tracking the gradual increase in the behavioral preference for humidity (Figure 1B). Indeed, a majority of log_2_(FC) values for 6 hours were between 30% and 70% of those measured at 12 hours (Figure 2G). In contrast, expression of *Sod3* (*superoxide dismutase 3*) remained stable until 6 h but had decreased by 12 h (Figure 2F, bottom left) consistent with its suppression perhaps participating in the metabolic or stress response to severe dehydration. Quenching thirst tended to return some transcript levels towards their sated baseline, although with different magnitude for different genes and clusters. However, transcript levels of many genes continued to increase after rehydration (Figure 2F, bottom right, Figure 2G), suggesting a prolonged action beyond the rehydration period. Together these results illustrate a diversity of water-deprivation induced gene regulation patterns.

### Astrocytic *aay* bidirectionally regulates water consumption

We tested whether glial genes identified in our scRNA-seq analysis regulated water consumption in the CAFE assay (Ja et al., 2007)(Figures 3A and 3B). We targeted temporally-restricted RNAi expression to adult glia using Repo-GAL4 in combination with a ubiquitously expressed temperature-sensitive GAL80 (McGuire et al., 2003)(Figure 3C). The strongest change in water consumption was produced with *astray* (*aay*). Adult-restricted RNAi-mediated knock down of *aay* reduced water consumption whereas overexpression increased it (Figures 3B, 3C, S3D and S3E). Notably, *aay* appears to specifically regulate water consumption, as food consumption was unaffected by *aay* RNAi (Figure S3F).

**Figure 3.**
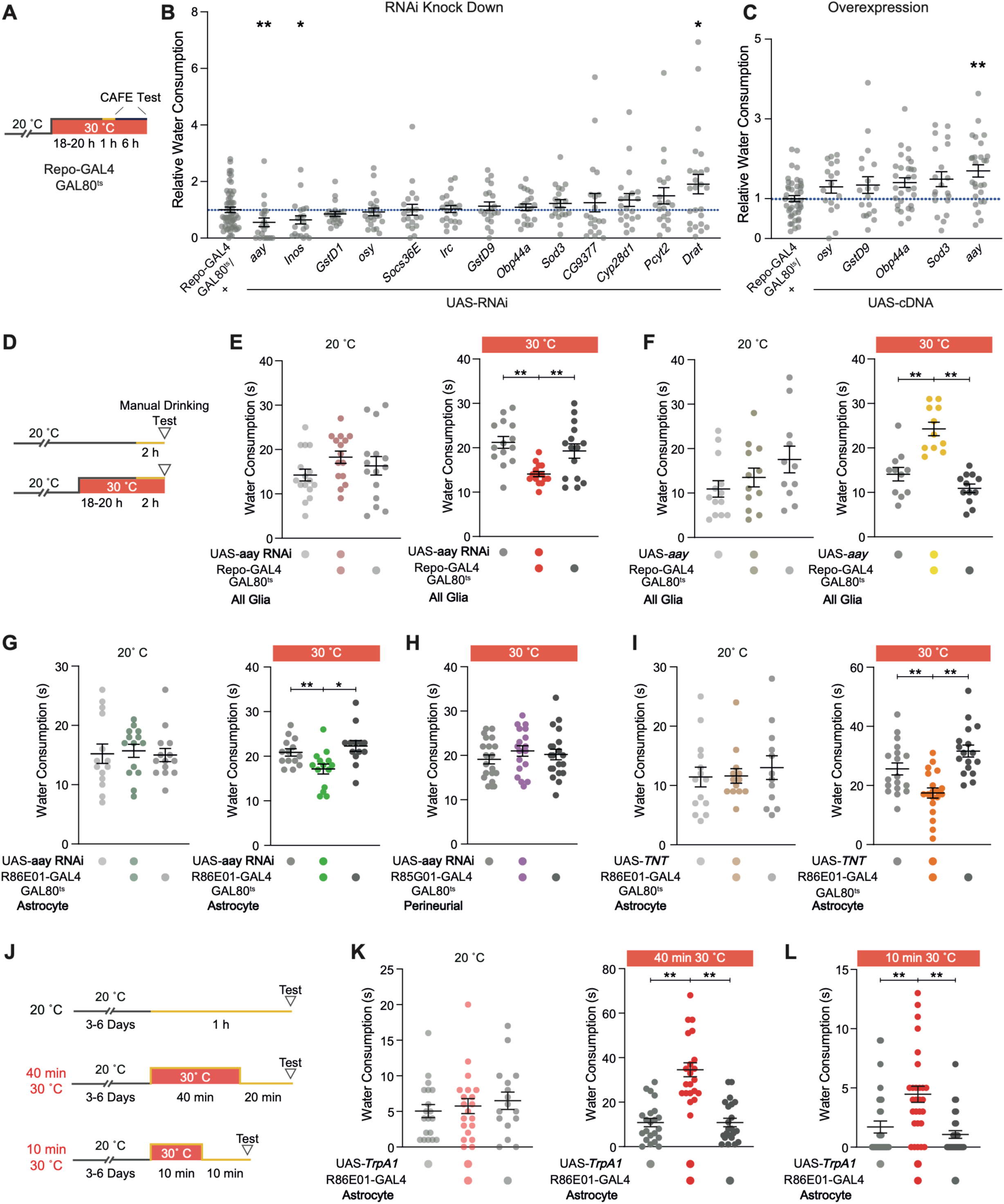
Astrocytic *aay* is a novel regulator of water consumption. (**A**) Schematic for temperature control of GAL80^ts^/GAL4 driven expression of UAS-RNAi or UAS-cDNA transgenes with CAFE test. Orange section of line indicates period of water restriction. (**B**) Water consumption in the CAFE assay of flies with RNAi knock down of candidate genes. Dotted blue line indicates normalization to control Repo-GAL4 flies, equal to 1. * p<0.05, ** p<0.01 two-tailed Mann-Whitney test n_RNAi_ = 20-25, n_Repo-GAL4_ = 50. (**C**) Water consumption in CAFE for UAS overexpression of targets. ** p<0.01 two-tailed Mann-Whitney test n_RNAi_ = 17-29, n_Repo-GAL4_ = 43. (**D**) Schematic for temperature control of GAL80^ts^/GAL4 driven UAS-RNAi or UAS-cDNA transgenes with manual water feeding assay. Orange section of line indicates period of water restriction. (**E**) RNAi knockdown of *aay* reduces water consumption (n = 14-16). (**F**) Overexpression of *aay* increases water consumption (n = 11-13). (**G**) Astrocyte-specific RNAi knockdown of *aay* reduces water consumption (n = 13,14). (**H**) RNAi knockdown of *aay* in perineurial glia does not alter water consumption (n = 18-20). (**I**) Preventing vesicular transmission with tetanus-toxin (TNT) expression in astrocytes reduces water consumption (n = 12-19). (**J**) Temperature regimen for TrpA1 activation of astrocytes in panels (K) & (L). Orange section of line indicates period of water restriction. (**K**) Astrocyte activation for 40 min increases water consumption (n = 16-22). ** p<0.01, Kruskal-Wallis ANOVA with Dunn’s multiple comparisons test. (**L**) Astrocyte activation for 10 min increases water consumption (n = 28-30). ** p<0.01, Kruskal-Wallis ANOVA with Dunn’s multiple comparisons test. Normality was assessed using Shapiro-Wilk test. * p<0.05, ** p<0.01, *** p<0.001, **** p<0.0001 Ordinary one-way ANOVA with Dunnett’s multiple comparisons test, unless otherwise stated. Individual data points are single flies. Data are mean +/-Standard Error of the Mean (SEM). See also **Figure S3**.

We noticed a significant number of deaths in the CAFE assay when manipulating *aay* (Figures S3G, S4D and S4E). This could result from sensitivity to the stress of dehydration, or idiosyncrasies of the assay itself (Park et al., 2018). To circumvent this issue we employed a manual water consumption assay that is less challenging for the animals (Jourjine et al., 2016). These experiments revealed similar results to the CAFE assay when knocking down or overexpressing *aay* in all glia (Figures 3D-3F). We next tested whether *aay*-induced changes in water consumption could be assigned to a specific glial cell-type (Kremer et al., 2017). Expressing *aay* RNAi in astrocytes, but not in perineurial glia, reduced water consumption indicating the specific importance of astrocytic *aay* in controlling drinking (Figures 3G and 3H).

The *aay* gene encodes a phosphoserine phosphatase which converts phosphoserine into D- and L-serine (Prokopenko et al., 2000; Salzberg et al., 1994; Wood et al., 1996). In mammals, the gliotransmitter D-serine is packaged into astrocytic vesicles and is released in a SNARE and calcium-dependent manner (Kang et al., 2013; Martineau et al., 2008; Mothet et al., 2005; Mustafa et al., 2004; Schell et al., 1997). *Drosophila* astrocytes express vesicular machinery including synaptobrevin (*Syb*) and Syntaxin 1A (*Syx1A*) (Figure S3M and S3N). To test whether vesicular release from astrocytes is necessary for water consumption, we expressed a tetanus-toxin (TetX) transgene using an astrocyte-specific GAL4. Tetx abolishes evoked release by cleaving the vesicle-associated membrane protein synaptobrevin. Temporally-restricted expression of TetX in astrocytes reduced water consumption (Figure 3I) (Kremer et al., 2017).

We also tested whether evoking Ca^2+^ entry into astrocytes promoted water consumption by expressing a transgene encoding the Ca^2+^ permeable temperature-sensitive TrpA1 channel specifically in astrocytes (*R86E01*>*TrpA1*). We first tested flies with a 2 h ‘fictive dehydration’ step at the TrpA1 activation temperature of 30 °C; however, this manipulation reversibly paralyzed *R86E01*>*TrpA1* flies, as previously described (Zhang et al., 2017). We therefore shortened the 30 °C heat stimulation to 40 min and allowed flies to recover at 20 °C for 20 min (Figure 3J). This procedure substantially increased water consumption compared to controls carrying only the *R86E01-GAL4* or UAS-*TrpA1* transgenes or those left at 20 °C for the whole procedure (Figure 3K). Since vesicular release occurs promptly following astrocytic activation (Kang et al., 2013), we also further restricted the heat incubation to 10 min. This brief astrocyte activation was also sufficient to increase water consumption (Figures 3L and S3L). Together these data indicate that *aay* expression and vesicular release from activated astrocytes regulate water consumption.

### D-serine facilitates water consumption via NMDA receptors

To further assess the role of D-serine as a gliotransmitter involved in water consumption, we supplemented the flies’ diet with D-serine and measured water intake (Figures 4A, 4B and S4A). Feeding flies with D- but not L-serine increased water consumption (Figure 4C). D-serine is known to be an enantiomer-specific co-agonist for NMDA-type glutamate receptors (NMDARs) (Kleckner and Dingledine, 1988; Mothet et al., 2000). We therefore tested if *aay*-dependent drinking defects could be rescued by dietary supplementation with D-serine. Glial expression of *aay* RNAi was induced in flies previously fed with medium supplemented with D-serine (Figure 4B). D-serine feeding was sufficient to restore the *aay* induced drinking deficit to a normal level (Figure 4D). Control experiments showed that flies have no preference for D-serine containing food, and that D-serine does not alter food consumption (Figures S4B and S4C).

**Figure 4.**
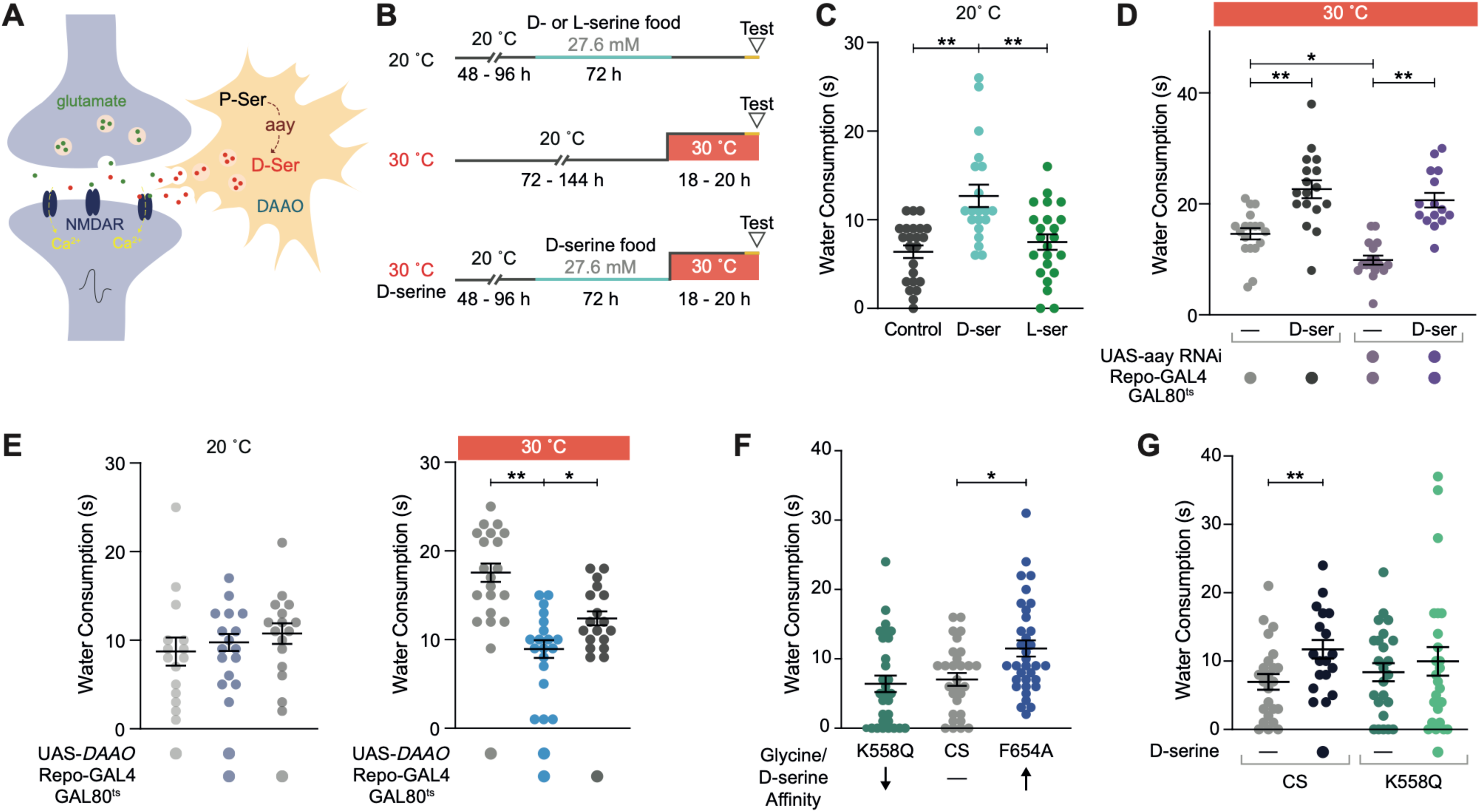
D-serine modulates water consumption via NMDARs. (**A**) Model of tripartite glutamatergic synapse showing *aay*-dependent synthesis of D-serine in astrocytes. (**B**) Protocol for D-serine feeding and RNAi induction with water consumption experiments (C) – (G). Light blue section of lines indicate time on D- or L-serine food. Orange section of lines indicate period of water restriction. (**C**) D- but not L-serine feeding increases water consumption (n = 20-23). ** p<0.01, Kruskal-Wallis ANOVA with Dunn’s multiple comparisons test. (**D**) D-serine feeding rescues the water consumption defect in flies with *aay* knockdown (n = 15-18). * p<0.05, **p<0.01, Ordinary one-way ANOVA with Dunnett’s multiple comparisons test. (**E**) Glial overexpression of D-amino acid oxidase (*DAAO*) reduces water consumption (n = 16-20). * p<0.05, ** p<0.01, Ordinary one-way ANOVA with Dunnett’s multiple comparisons test. (**F**) Flies harboring the F654A single site mutation in NMDAR1 exhibit increased water consumption. This mutation increases affinity for glycine/D-serine. (n = 29-34). * p<0.05, Kruskal-Wallis ANOVA with Dunn’s multiple comparisons test. (**G**) Flies harboring the K558Q amino acid substitution in NMDAR1 are insensitive to the D-serine induced increase in water consumption (n = 18-25). ** p<0.01, Two-tailed Mann-Whitney test. Individual data points are single flies. Data are mean +/- SEM. See also **Figure S4.**

D-serine is degraded by D-amino acid oxidase (DAAO), which is encoded by the *CG12338* and *CG11236* genes in *Drosophila* (Dai et al., 2019; Pollegioni et al., 2007). Since *CG11236* expression is barely detectable in the brain we focused on *CG12338* (Dai et al., 2019). Overexpressing a transgene for the *CG12338* DAAO in glia suppressed water consumption (Figure 4E). Taken together, the *aay* manipulation, dietary supplementation and DAAO data suggest that D-serine can bi-directionally regulate water consumption.

NMDARs are voltage and ligand-gated ion channels. Efficient channel activation requires binding of L-glutamate to the NR2 subunit and a co-agonist glycine or D-serine, which bind to the same site on the NR1 subunit (Laube et al., 1997). Prior neuronal depolarization is also required to remove a Mg^2+^ block allowing ligand-gated channel conductance to occur. We therefore tested whether D-serine facilitated water consumption via NMDAR activation. To address this, we used flies harboring single-site mutations (K558Q and F654A) in the *NMDAR1* gene which encodes the fly NR1 subunit. These amino acid substitutions are equivalent to K544Q and F639A in the mammalian NR1 subunit (Miller, 2004; Troutwine et al., 2019; Yoneda et al., 1993). Glycine affinity is increased in receptors bearing F639A/F654A and decreased in those with K544Q/K558Q, whereas glutamate affinity is unaltered (Dickinson et al., 2007; Wafford et al., 1995). Measuring innate water consumption revealed that flies harboring the F654A substitution in NR1 drank more than controls (Figure 4F). In addition, K558Q flies exhibited normal levels of water consumption but they were unable to acquire the D-serine induced increase in drinking (Figure 4G). These data demonstrate that NMDAR co-agonist affinity directly impacts water consumption.

### D-serine is a co-agonist of the *Drosophila* NMDA receptor

A prior study has demonstrated that D- or L-Serine feeding can regulate sleep in flies, and that this effect depends on NMDARs (Dai et al., 2019). However, it is currently an assumption that D-serine acts as a co-agonist of *Drosophila* NMDARs, as it does in mammals. Since NMDAR activation elicits increases in intracellular Ca^2+^, we expressed GCaMP7f broadly in NR1 positive (NR1+) neurons and performed *in-vivo* calcium imaging in the brain of head fixed flies while bath applying glycine, D-serine, or L-serine (Figure S5A and S5B). We chose to record from NR1+ neurons (*nmdar1-KIGAL4>GCaMP7f*) in the pars intercerebralis (PI) because they are easily accessed and identified, and therefore permitted reproducible measurements between flies (Figure 5A) (Dai et al., 2019). In addition, we measured neuronal responses in both Mg^2+^ free (0 mM Mg^2+^) and physiologically relevant concentrations of Mg^2+^ (4 mM Mg^2+^) (Figure 5B) to assess the importance of removing the Mg^2+^ occlusion from the channel pore (Ascher and Nowak, 1987; Mori et al., 1992). Bath application of NMDA with D-, but not L-, serine evoked robust activation of PI NR1+ neurons (Figures 5C and 5D). NMDA and glycine also produced an excitatory response but only in Mg^2+^ free conditions (Figure 5D). To validate that D-serine activation was occurring via NMDARs we co-applied the non-competitive NMDAR antagonist ketamine with NMDA and D-serine (Figure 5E) (Martin and Lodge, 1985). Inclusion of ketamine abrogated PI neuron responses to D-serine/NMDA application (Figures 5F, 5G and S5H). These data demonstrate that fly NMDARs are activated by D-serine.

**Figure 5.**
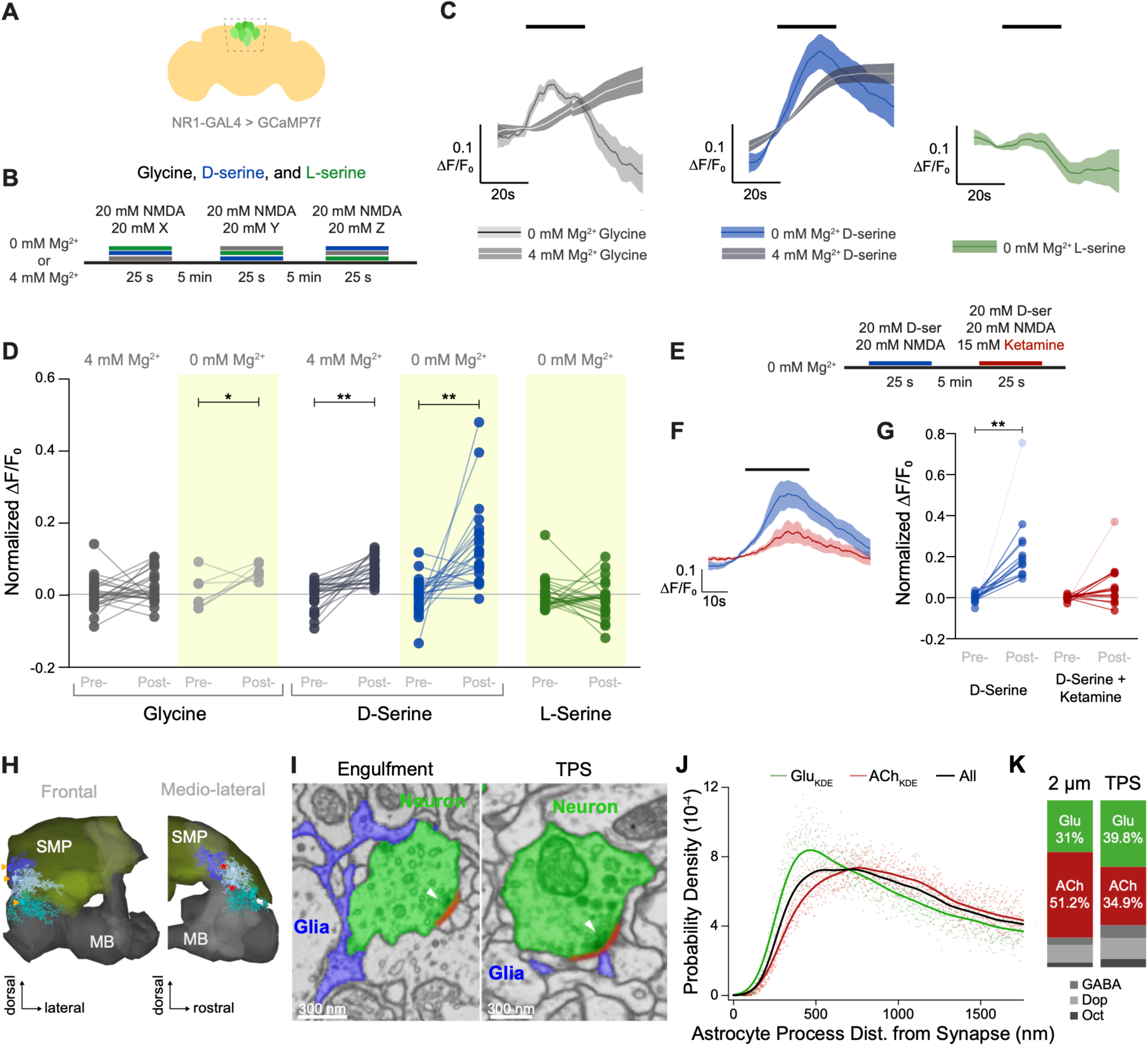
Astrocytes form tripartite synapses and D-serine is a co-agonist for NMDARs. (**A**). Illustration of imaging window for recording of Ca^2+^ responses in NMDAR1 expressing PI neurons. (**B**) Protocol of drug application for panels (C) & (D). Order of application was randomized for each fly. (**C**) Average traces for glycine, D-serine, and L-serine application in 0 or 4mM Mg^2+^ (n_gly_ = 8, 25; n_D-ser_ = 25, 26; n_L-ser_ = 28, respectively). Line is a moving average and shaded band is SEM. (**D**) D-serine, but not L-serine, activates NR1+ PI neurons (from left to right, n = 25, 25, 8, 8, 25, 25, 23, 23, 28, 28). * p<0.05, ** p<0.001, Kruskal-Wallis ANOVA with Dunn’s multiple comparisons test. (**E**) Protocol for ketamine application with NMDA, TTX, and D-serine. (**F**) Averaged traces for D-serine and NMDA (blue) and D-serine, NMDA, and ketamine (red) (n = 15). (**G**) Ketamine inhibits D-serine induced activation of NR1+ PI neurons. ** p<0.01, Kruskal-Wallis ANOVA with Dunn’s multiple comparisons test. Transparent dots indicate outliers that are > 2 SD from the mean. When outliers are excluded from the analysis the relationship still holds (see Figure S5F). (**H**) Astrocytes tile the SMP. 3D representation of three astrocytes (shades of blue) reconstructed from EM data shown with the mushroom body neuropil (grey) and the SMP neuropil (yellow), shown in a frontal and lateral view. Cell bodies are located outside the neuropil (yellow arrow heads), and processes in the neuropil have little overlap (*). (**I**) Astrocytes engulf synaptic boutons and contribute processes to tripartite synapses (TPS). Greyscale EM data with labeled glutamatergic boutons (green) and glial processes (blue). Presynaptic densities (arrow) and synaptic cleft (red) are indicated. Left: Example where glial processes contact a large proportion of a bouton’s membrane but not the synapse. Right: Example of a glial process directly adjacent to the synaptic cleft and opposite the presynaptic density (white arrow head). (**J**) Astrocytic processes are significantly closer to glutamatergic than cholinergic synapses in the SMP. Kernel density estimates (lines) of the probability distributions (points) of distances of Glu or Ach synapses or a random draw from both sets to 3 SMP based astrocytes. Only synapses in direct vicinity (2µm radius around processes <600nm thick) are considered. Statistical analyses shown in Figure S5. (**K**) Glutamatergic synapses are overrepresented in tripartite synapses vs. their representation in synapses in the general 2 µm vicinity of astrocytic processes. Ratios of synapses by predicted neurotransmitter usage are shown. See also Figure S5 and Video S1 for anatomy of astrocyte processes engulfing a presynapse and contributing to TPS.

### Astrocytes contribute to tripartite synapses in the *Drosophila* brain

Processes from mammalian astrocytes infiltrate the synaptic cleft, forming tripartite synapses (TPS) (Ventura and Harris, 1999). Proximity to the synapse allows for glia to participate in neurotransmission through uptake and recycling of neurotransmitters, and by release of gliotransmitters (Santello et al., 2012). A tripartite structure has been observed for *Drosophila* adult neuromuscular synapses, but the detailed anatomy of astrocytes in the adult central nervous system (CNS) is currently unclear (Danjo et al., 2011). To address this we analyzed astrocytes in the Superior Medial Protocerebrum (SMP) neuropil within the FlyWire project (flywire.ai) (Dorkenwald et al., 2022) which utilizes the Full Adult Female Brain (FAFB) transmission electron microscope dataset (Zheng et al., 2018). We fully reconstructed the structure of 11 astrocytes and four were extensively reviewed in order to reconstruct the finest tips of their processes. We found astrocytes to tile the neuropil and to be closely associated with trachea, as previously reported in the larval VNC and brain (Ma and Freeman, 2020; Pereanu et al., 2007; Stork et al., 2014) (Figures 5H, S5J and S5K). We also identified astrocytic processes that engulfed synaptic boutons and that contacted synaptic clefts like processes from postsynaptic neurons – the latter providing evidence of canonical tripartite synapses in the adult CNS (Figures 5I and S5L). In addition, astrocytes also often contact the postsynaptic neuron within 500 nm of the synaptic cleft.

To determine whether astrocytes associate with a particular class of synapse we used the prior computational predictions of neurotransmitter identity for neurons in the FAFB volume (Eckstein et al., 2020). We composed a vicinity profile by measuring distances of synapses from the fine processes of the 3 extensively reviewed SMP astrocytes and plotted their distance distributions categorized by presynaptic neurotransmitter. This analysis revealed that these astrocytic processes are significantly closer to glutamatergic (962nm ± 2nm SD) and GABAergic (971nm ± 3 nm) than cholinergic synapses (1061nm ± 1 nm) (Figures 5J, 5K, S5M and S5N). Astrocytes are therefore ideally positioned to modulate glutamatergic synapses in the adult fly brain.

### Astrocytes show heterogeneous responses to neurotransmitters

Our demonstration that acute astrocyte activation was sufficient to drive increases in water consumption, and their placement close to synapses, led us to hypothesize that astrocytes are active participants in regulating water consumption. Astrocytes have been proposed to facilitate fast synaptic activity by releasing gliotransmitters such as glutamate, ATP, and D-serine (Savtchouk and Volterra, 2018). In mammals, glutamate binding to mGluR5 triggers astrocytic D-serine release (Mustafa et al., 2009) and the concomitant presence of glutamate and D-Serine co-agonist is crucial for maximal NMDAR activation. We therefore tested whether fly astrocytes were responsive to glutamate application by recording *in vivo* astrocytic Ca^2+^ responses (*R86E01-GAL4*>*UAS-GCaMP7f*) (Figures 6A and 6B). We included 1 µM tetrodotoxin (TTX) in the bath to block voltage-gated sodium channels and thereby inhibit polysynaptic neurotransmission. Surprisingly, we were able to classify astrocytes within individual flies depending on whether bath applied glutamate evoked an increase, decrease, or no change in GCaMP fluorescence (Figure 6B). This unexpected result led us to determine if a similar array of responses could be observed with other neurotransmitters. We used acetylcholine (ACh) because it is the predominant excitatory neurotransmitter in the fly brain, and ATP because it triggers a robust response in mammalian astrocytes (Guthrie et al., 1999; Lee and O’Dowd, 1999). Differential astrocytic responses to ACh and ATP were also evident (Figures 6C-6F). Moreover, the same astrocytes exhibited similar responses to the different neurotransmitters. For instance, a substantial proportion of astrocytes were activated both by glutamate and ATP (Figures 6G, 6H, S6A and S6B) and a substantial proportion of them had matched responses to all three neurotransmitters (Figure 6I).

**Figure 6.**
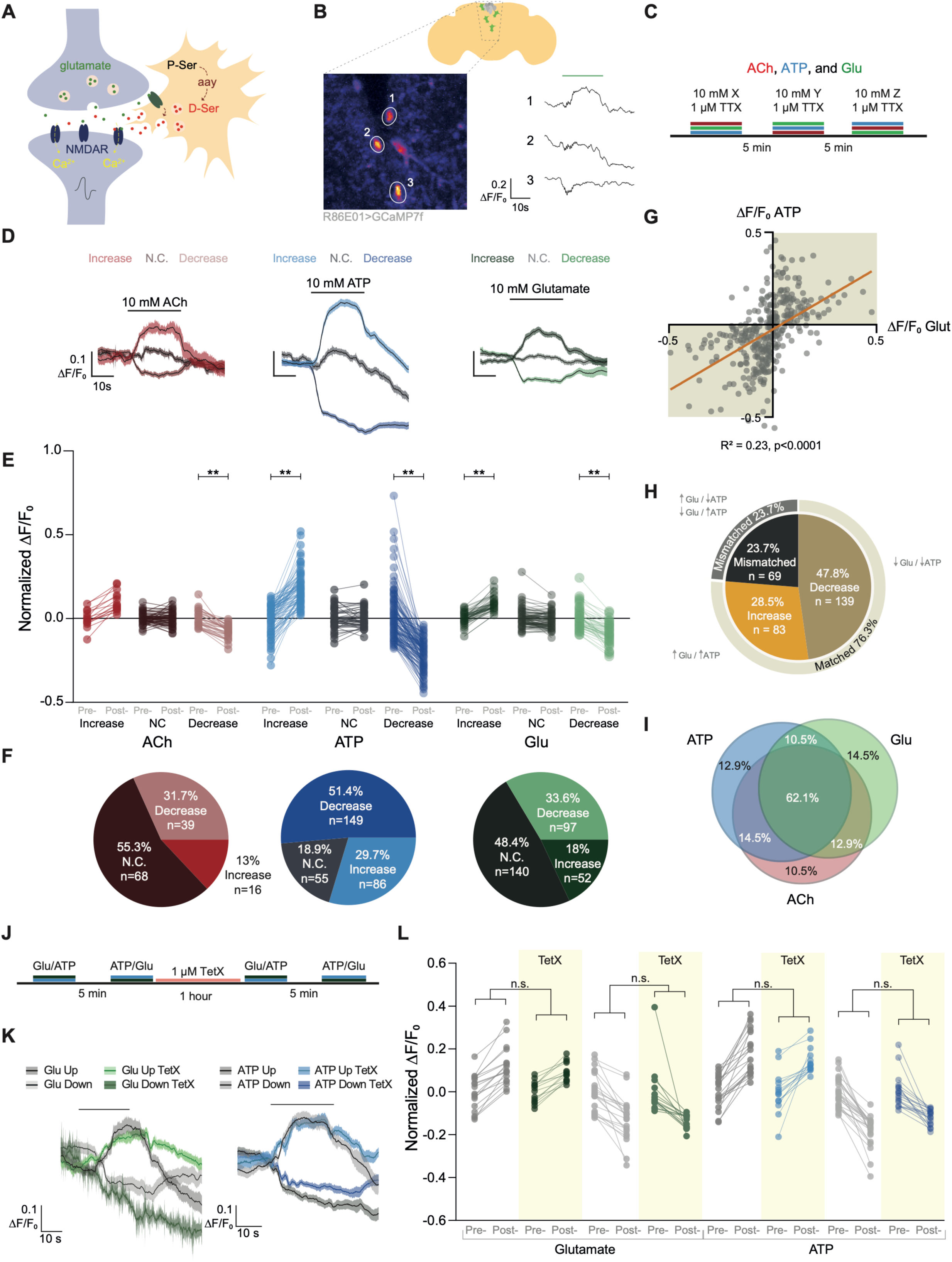
Astrocytes are differentially responsive to neurotransmitters. (**A**) Illustration of possible glutamate-evoked D-serine release from astrocytes. (**B**) Astrocytes adjacent to the pars intercerebralis show differential responses to glutamate application (green bar). (**C**) Protocol for drug application with randomized order for Acetylcholine (ACh), ATP, and Glutamate (Glu). (**D**) Average traces for ACh, ATP, and glutamate partitioned by the type of response (activated, no change, or inhibited). Responses were determined as follows: activated if µ_post-drug_ > σ_pre-drug_ + µ_pre-drug_, no change if µ_post-drug_ fell within σ_pre-drug_ + µ_pre-drug_ and inhibited if µ_post-drug_ < σ_pre-drug_ - µ_pre-drug_. Line is smoothed average and shaded band is SEM. (**E**) Neurotransmitters induce both excitatory and inhibitory responses in astrocytes. Paired datapoints represent responses from a single astrocyte and group is assembled from multiple flies (n_Ach_ = 16, 68, 39; n_ATP_ = 86, 55, 149; n_glut_ = 52, 140, 97). ** p<0.01, Kruskal-Wallis ANOVA with Dunn’s multiple comparisons test. (**F**) Proportions of astrocytes sorted by response direction following acetylcholine, ATP, and glutamate application. (**G**) Average ΔF/F_0_ during drug application for glutamate plotted against ΔF/F_0_ for ATP shows most astrocytes that are excited by glutamate are also excited by ATP and vice versa. (**H**) Astrocytes are more likely to have matched responses between glutamate and ATP. Chi-square test between matched vs. mismatched astrocytes. Fisher’s exact test matched vs. mismatched p = 2.35e-11, OR = 3.24. (**I**) Venn diagram showing considerable overlap of matched responses of astrocytes to all three neurotransmitters. (**J**) Protocol for tetanus toxin (TetX) application. (**K**) Average traces for glutamate and ATP evoked excitatory and inhibitory responses with and without TetX. Line is smoothed average and shaded band SEM. (**L**) Blocking vesicular transmission does not suppress astrocyte responses to transmitter application (left to right, n_Pre-TetX ATP_ = 21, 6; n_Post-TetX ATP_ = 16, 35; n_Pre-TetX Glut_ = 18, 11; n_Post-TetX Glut_ = 14, 9). n.s. > 0.05 Kruskal-Wallis ANOVA with Dunn’s multiple comparisons test. Individual data points are single astrocytes across multiple animals. See also **Figure S6**.

We also used TetX to eliminate all vesicular release and rule out the possibility that evoking local monosynaptic transmission accounts for the differential astrocytic responses (Figure S6C) (Yamasaki et al., 1994). Although there was no change in the magnitude of astrocyte responses to either ATP or glutamate following a 1 h TetX application, there was a slight reduction in the percentage of ATP responsive astrocytes (Figures 6J-6L and S6F). We verified that the TetX was effective by imaging a polysynaptic odor response in mushroom body output neuron (MBON)-γ5β′2a. TetX application eliminated odor-evoked calcium responses in this neuron (Figures S6D and S6E). Together, these results demonstrate that neurotransmitters can evoke direct stereotyped activation or inhibition of astrocytes. Differential astrocytic responses to neuronal activation were previously observed in the larval ventral nerve cord where astrocytes displayed either a sustained inward current, or a brief inward followed by a longer outward current (MacNamee et al., 2016).

### Water deprivation modulates functional properties of astrocytes

Having characterized astrocytic responses to neurotransmitters, we tested whether these properties changed in water deprived flies. We performed *in vivo* Ca^2+^ imaging in sated, dehydrated, and food deprived flies (Figure 7A). To prevent deprived flies from being ‘re-satiated’ by the application of normal extracellular saline, we adjusted the saline composition to reflect the desired physiological state of the animal (Cheriyamkunnel et al., 2021; Jourjine et al., 2016). Astrocytes did not show an obvious Ca^2+^ response to a brief increase in osmolarity (Figure 7B). Similar to sated flies, water and food deprived flies showed differential astrocytic responses to glutamate and ATP (Figures 7C-7J and S7A-S7C). However, only water deprived flies showed an increase in the proportion of astrocytes that were excited by glutamate, while the proportion of ATP responses remained the same (Figures 7F and 7J). Glutamate and ATP excitatory responses also appeared to be prolonged in thirsty flies compared to controls (Figures 7K, 7L, and S7D). Finally, compared to sated flies, thirsty flies had a higher proportion of mismatched astrocyte responses to ATP and glutamate, suggesting a loss of correlated responses (Figure S7G). In sum, water deprivation increases the proportion of astrocytes that are activated by glutamate and sustains glutamate-evoked Ca^2+^ responses.

**Figure 7.**
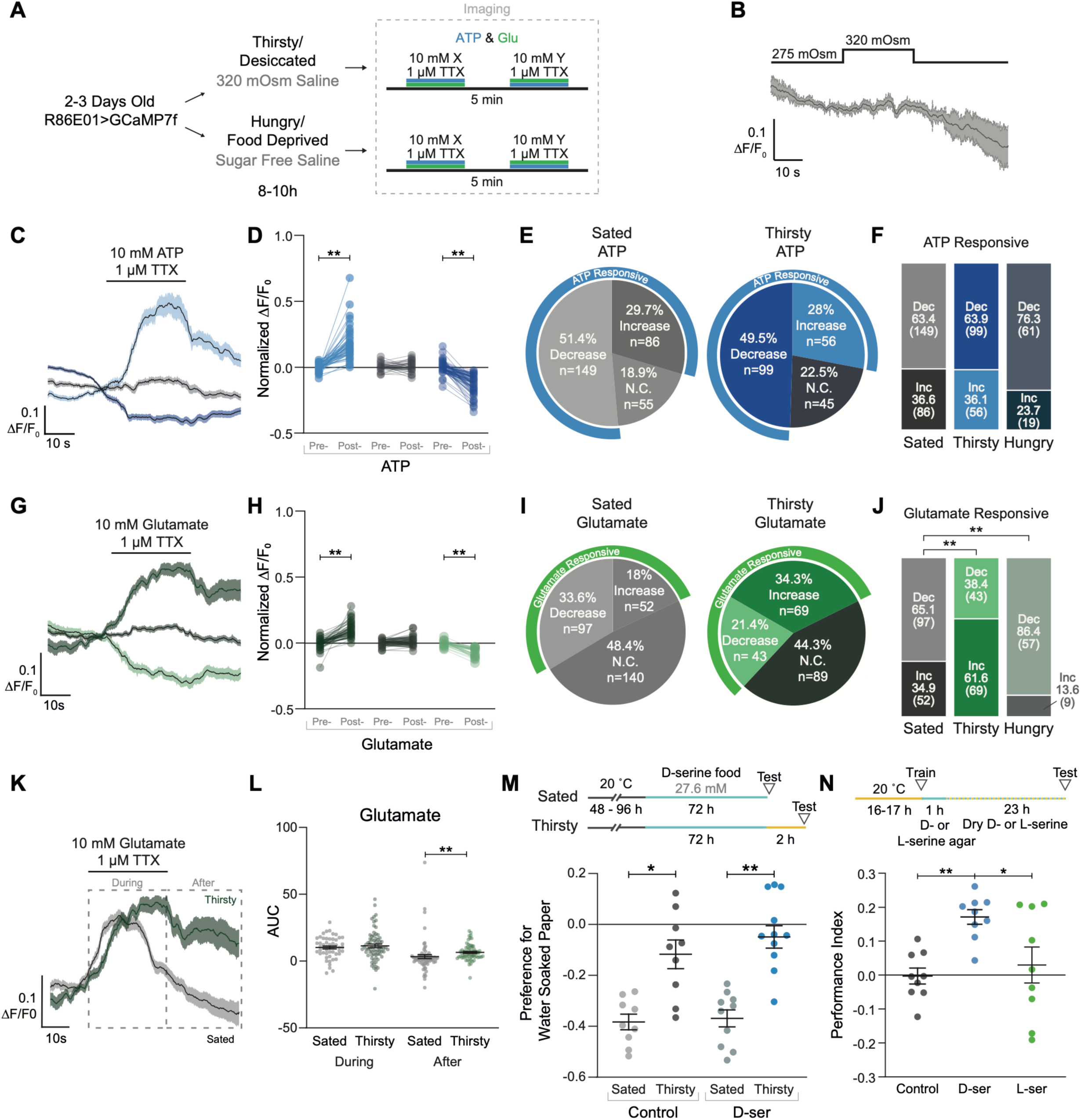
Water deprivation changes physiological properties of astrocytes. (**A**) Protocol for water and food deprivation with calcium imaging experiments. (**B**) Acute exposure of high osmolarity saline does not induce a calcium response in astrocytes. (**C**) Average traces for ATP partitioned by response type in thirsty flies. Line is smoothed average and shaded band is SEM. (**D**) Responses Pre- and Post- treatment of ATP separated by categorical responses in water deprived flies. ** p<0.01, Kruskal-Wallis ANOVA with Dunn’s multiple comparisons test. (**E**) Proportion of astrocyte responses to ATP application in sated and thirsty flies. (**F**) Astrocyte responses to ATP do not change with deprivation states. n.s > 0.05 Fisher’s exact test with Bonferroni correction. (**G**) Average traces for glutamate partitioned by response type in thirsty flies. Line is smoothed average and shaded band SEM. (**H**) Responses Pre- and Post- treatment of glutamate separated by categorical responses in water deprived flies. ** p<0.01, Kruskal-Wallis ANOVA with Dunn’s multiple comparisons test. (**I**) Proportions of astrocyte responses type to glutamate application in sated and thirsty flies. (**J**) Deprivation states differentially regulate astrocyte responsivity to glutamate application. The number of glutamate responsive astrocytes increases in thirsty flies, and decreases in hungry flies. ** p<0.01, Fisher’s exact test with Bonferroni correction. (**K**) Excitatory astrocyte responses to glutamate application are prolonged in thirsty animals. (**L**) Area Under the Curve (AUC) of sections of traces marked in (K). ** p<0.01, Kruskal-Wallis ANOVA with Dunn’s multiple comparisons test. (**M**) Flies pre-fed D-serine do not show differences in innate water seeking in the T-maze. * p<0.05, ** p<0.01, Kruskal-Wallis ANOVA with Dunn’s multiple comparisons test. (**N**) Flies fed D-serine between training and testing show increased water memory performance. * p<0.05, ** p<0.01, Kruskal-Wallis ANOVA with Dunn’s multiple comparisons test. For *D, H and L) individual data points are single astrocytes across multiple animals. See also **Figure S7**.

### D-serine promotes water-memory expression

Prior work has implicated specific neuronal pathways in regulating levels of water consumption and in the thirst-dependent control of innate and learned water-seeking (Jourjine et al., 2016; Lin et al., 2014; Landayan et al., 2021; Senapati et al., 2019). Given that astrocytes broadly innervate the neuropil we reasoned that D-serine might provide a more general signal promoting water procuring behaviors. We therefore tested whether D-serine could promote water vapor seeking and memory expression in water-sated flies, like it did for drinking. Interestingly, the same 3-day feeding of D-serine that increased drinking did not promote attraction to water vapor (Figure 7M). We next tested for an effect on the thirst-dependent expression of water memory performance. Water-deprived flies were trained to associate an odor with water reward and after training they were housed in vials with agarose for 1 h then dry vials containing either nothing, D-serine, or L-serine powder. Only flies fed with D-serine showed 24 h water memory performance (Figure 7N). These data demonstrate that D-serine promotes the expression of water-seeking memory, in addition to drinking behavior.

## Discussion

### Single-cell RNA-seq reveals a glial response to water-deprivation

Water deprivation reduces the volume and increases the osmolality of an animal’s hemolymph or blood. These changes induce multiple adaptations throughout the body, such as changes in the activity of osmosensory nuclei in the brain (the LT in mammals and ISNs in *Drosophila*) as well as production and release of neuropeptide or neurohormone modulators (Augustine et al., 2020; Gáliková et al., 2018; Jourjine et al., 2016; Landayan et al., 2021; Matsuda et al., 2020; Senapati et al., 2019; Zimmerman et al., 2017). These molecules modulate organs controlling water retention, excretion and blood pressure, and the activity of neuronal circuits. In *Drosophila* these circuits include specific reinforcing dopaminergic neurons (Hsu et al., 2020; Lin et al., 2014b; Senapati et al., 2019), that ultimately influence motivational control of water-seeking and drinking. Our single-cell analyses of water-deprived flies did not identify significant transcriptional changes within peptidergic or monoaminergic neurons in water-deprived flies, perhaps due to the relatively small numbers of these neurons and their representation in our datasets. Instead, our studies revealed astrocytes and other glia to be the principal transcriptionally responsive cells in the fly brain.

Although the glial expression of many genes changed in thirsty flies, our initial loss-of-function RNAi screen only revealed potential roles for *aay*, *inos* (myo-inositol-1-phosphate synthase) and *Drat* (Death resistor Adh domain containing target) in the control of drinking behavior. Results for *inos* and *Drat* were not corroborated by additional testing. Nevertheless, we note that induction of myo-inositol synthesis and uptake is a known cellular response to osmotic stress (Sacchi et al., 2013). *Drat* might also be involved in the resistance to the stress of dehydration (Chen et al., 2012). We similarly speculate that many of the other thirst-regulated genes play roles in maintaining glial function under stress, without necessarily impacting the fly’s fluid intake in a measurable way.

### Glial D-serine is required for motivated water procuring behaviors

Since the *aay*-encoded phosphoserine phosphatase is required to synthesize serine we hypothesized that *aay* flies might be deficient in motivated thirst-dependent behavior because they lacked the gliotransmitter D-serine. Whereas glial *aay* loss-of-function decreased drinking, glial *aay* overexpression enhanced water consumption. A key approach to verifying a role for D-serine was the ability to alter fly behavior directly by providing D-serine in the diet. Dietary D- but not L-serine restored the drinking defect of *aay*-deficient flies. Moreover, feeding D- but not L-serine to water-sated flies mimicked the behavioral effect of water-deprivation – promoting both drinking and water-seeking memory expression. We therefore conclude that gliotransmission of D-serine is an essential element of thirst-state directed behavioral modulation. Changes in hunger-directed feeding behavior were not evident. At present we do not know if dietary D-serine floods the entire fly and bypasses the need for glial release. Interestingly, whereas our feeding experiments demonstrated D-serine enantiomer specific *aay* rescue and gain of function effects on thirst-related behaviors, a previous study (Dai et al., 2019) rescued a fly sleep defect with both L- and D-serine feeding. However, rescue of sleep was dependent on intestinal serine racemase conversion of L- to D-serine. Since the authors also identified intestinal NMDAR1 expressing neurons (Dai et al., 2019), we speculate that L-serine conversion in the gut produces enough D-serine to alter sleep via that route, but it produces insufficient quantities to act in the brain (Tomita et al., 2015; and our studies here).

### Does water-deprivation prime astrocytes to release D-serine?

We noted that increased *aay* expression resulted in detection of a larger proportion of astrocytes (74% versus 56%) expressing *aay* following water deprivation. This potential broadening of expression might indicate that water deprivation induces *aay* to meet a forthcoming demand for D-serine release, and/ or to replenish it following release from these astrocytes. Interestingly, a similar number of astrocytes (34% versus 18%) also gained sensitivity to exogenously applied glutamate in thirsty flies. Although we do not know if the same astrocytes are involved, it seems plausible that thirst increases both astrocytic expression of *aay* and the number of glutamate responsive astrocytes, so that glutamate-releasing neurons can evoke their D-serine release. Although our loss-of-function experiments reveal the importance of astrocytic *aay*, induction was also evident in cortex, ensheathing, and surface glia. The role of *aay* induction in these other glial types is currently unclear. Elevated expression of *aay* was attenuated when flies were permitted to quench their thirst, consistent with induction serving an enhanced need for D-serine synthesis.

### Astrocytic D-serine facilitates the activity of glutamatergic circuits

We established that D-serine functions as an NMDAR co-agonist in *Drosophila,* like it does in mammals (Mothet et al., 2000). Neuronal NMDA-evoked Ca^2+^ transients were potentiated by D-serine co-application and blocked by ketamine. Moreover, feeding D-serine did not promote drinking in flies harboring a K558Q point substitution in the NMDAR1 subunit, that renders it insensitive to D-serine (Troutwine et al., 2019). Importantly, our ultrastructural analyses identified typical tripartite synapses within the adult fly brain, as well as a new and more common anatomical motif where astrocytic processes engulf presynaptic boutons. Astrocytic processes are therefore well placed to potentially support synaptic transmission acting on either the pre- or post-synaptic neuron. Moreover, astrocytic processes in the SMP appear to be preferentially associated with glutamatergic synapses, despite cholinergic synapses being the most abundant in this region. We therefore propose that thirst-regulated astrocyte-released D-serine facilitates NMDA-type currents so that the relevant glutamatergic synapses undergo short-term potentiation, which allows these water-procuring neural circuits to function more efficiently. If D-serine release itself depends on glutamate-evoked astrocyte activity at active tripartite synapses then its use will be need-dependent and circuit-specific. It is also conceivable that astrocytic processes at tripartite synapses can be remodeled by osmosensitive swelling and retraction (Wang and Parpura, 2018). Further work will be required to locate the critical D-serine modulated connections and circuits in the fly brain. We assume different circuits will underlie D-serine regulated learned water seeking and consumption (Hsu et al., 2020; Jourjine et al., 2016; Landayan et al., 2021; Lin et al., 2014b; Senapati et al., 2019).

### Do glia directly sense water deprivation?

Although glia are ideally positioned for both integrating hemolymph metabolic and osmotic states and instructing neuronal activity we did not observe an obvious astrocytic Ca^2+^ response when the osmolarity of the extracellular solution was briefly increased. However, it is possible that astrocyte activation requires a longer and more sustained duration of exposure to hyperosmotic solution, that they are instead sensitive to hypovolaemic changes, or that water deprivation does not induce a Ca^2+^ response in astrocytes (we also do not know if thirst-driven glial transcriptional responses are Ca^2+^-dependent). In addition, whereas fly perineurial and subperieurial glia provide the barrier between the hemolymph and the brain, cortex glia wrap neuronal cell bodies, and astrocytes permeate the neuropil and associate with synaptic compartments. It is therefore plausible that an interaction between the layers of glia permits transduction of osmosensing signals to astrocyes. For example, Ca^2+^ oscillations in cortex glia, which require the Na^+^/Ca^2+^, K^+^ exchanger zydeco (Melom and Littleton, 2013), could influence astrocyte activity/Ca^2+^, as they are directly connected (Farca Luna et al., 2017).

Previous work in mammals has implicated astrocytic Na_(X)_ channels in sodium sensing within the SFO. Na_(X)_ are coupled to Na^+^/K^+^-ATPases that potentiate metabolic activity facilitating astrocytic release of lactate, which stimulates adjacent GABAergic neurons to ultimately promote salt aversion in the mouse (Shimizu et al., 2007). Our single-cell sequencing data reveals glial-specific expression of a number of potential osmo-sensing molecules. Several solute carrier (SLC) transporters localize to glial clusters such as *Ncc69*, *NKCC* and *inebriated* (*ine*) (Data S2). Interestingly, the *ine* encoded Na^+^/Cl^-^ dependent neurotransmitter transporter is required for water homeostasis in the Malpighian tubules (Luan et al., 2015). Additionally, *ine* negatively regulates neuronal activity, which could be a mechanism by which astrocytes couple osmotic regulation with neurotransmission (Huang and Stern, 2002). Several TRP genes encode mechanosensitive channels that are responsive to changes in cell volume (Pedersen et al., 2011). The *pain* and *wtrw* TRP genes were exclusively detected in glia (Data S2). Finally, glia exhibit restricted expression of the inwardly rectifying potassium channel genes *Irk2 and Irk3*, which are necessary for proper fluid excretion and renal function in *Drosophila* (Evans et al., 2005). Taken together, the transcriptomes indicate glia have potential for sensing osmotic changes and regulating osmotic balance in the brain.

### Thirst specificity

Although in lab conditions *Drosophila* are raised on food that contains water, they can selectively seek either food or water when necessary, and prior work has established that they possess independent control mechanisms for food and water seeking (Hsu et al., 2020; Krashes et al., 2009; Landayan et al., 2021; Lin et al., 2014; Senapati et al., 2019). Thirst and hunger are however also coupled on a behavioral and neuronal level. For instance, prandial thirst is triggered by food ingestion to preemptively recalibrate blood osmolality (Augustine et al., 2020; Gutman and Krausz, 1969; Jourjine et al., 2016; Zimmerman et al., 2016). In this study, we found that D-serine apparently distinguishes between water and food consumption. In addition, astrocytes showed opposing physiological changes with food deprivation compared to water deprivation (Figure 7J), and expression of genes regulated by thirst barely changed when flies were hungry (Figure S2C). D-serine elevation, either by *aay* overexpression or D-serine feeding, promoted expression of water-seeking memory and water consummatory behaviors. Therefore *aay*-dependent D-serine signaling may provide a general thirst modulation, as opposed to only regulating seeking (Landayan et al., 2021) or drinking (Jourjine et al., 2016). Together, these results indicate that astrocytic D-serine represents a specific mechanism that acutely modulates water procurement. Further work is required to understand how thirst-driven D-serine interacts with other thirst-dependent modulatory signals. It will also be interesting to determine whether D-serine influences sodium appetite, and interacts with modulatory processes relevant to other motivational states (Cheriyamkunnel et al., 2021; Jung et al., 2020; McKinley et al., 2019; Senapati et al., 2019; Stricker and Hoffmann, 2007; Zhang et al., 2018).

### Could D-serine be a conserved state-mechanism in mammals?

Thirst has been recently described to alter brain-wide population dynamics in the mouse (Allen et al., 2019). Since astrocytes permeate all regions of the brain, it is possible that D-serine contributes to the broad thirst-driven modulation of neural activity. In addition, we note that a lower D-serine serum concentration has been linked to patients suffering from Schizophrenia (Hons et al., 2021), 6-20% of whom are polydipsic.

## Author contributions

Designed research A.P., V.C., N.O., D.A., C.T.D., E.M., S.W., Performed research A.P., V.C., N.O., D.A., C.T.D., E.M., Analyzed data A.P., V.C., N.O., D.A., C.T.D., D.S., S.W., Resources D.S. S.W. and V.C., Writing S.W., A.P., V.C., N.O., Supervision D.S., S.W. Funding Acquisition S.W., D.S.

## Acknowledgments

We thank Bernard Moussian, Yi Rao, Stefanie Schirmeir, the Vienna *Drosophila* Resource Center and Bloomington *Drosophila* Stock Center for flies. We are grateful to Hubert Slawinski at the Wellcome Trust Centre for Human Genetics for running single-cell transcriptomics and sequencing, Lucy Garner, Paul Brodersen and the Bioconductor community for assistance and advice with single-cell transcriptomics analysis, and Ruth Brain for lab assistance. We thank the Princeton FlyWire team, Murthy and Seung labs for development and maintenance of FlyWire (supported by BRAIN Initiative grant MH117815), and the Princeton EM proofreading team for contributing neuronal edits. We acknowledge Philipp Schlegel for help implementing navis-related code and Joseph Hsu for proof reading astrocyte morphology. We thank members of Waddell, Sims and Goodwin groups for discussion. A.P., V.C., C.D.T and S.W. were supported by an ERC Advanced Grant to S.W. (789274). V.C. was additionally supported by starter funds from Durham University, N.O was funded by Wellcome Collaborative Awards to S.W. (203261/Z/16/Z and 221300/Z/20/Z). D.A. was supported by a Wellcome Collaborative Award to D.S. and S.W. (209235/Z/17/Z). E.M. was funded by an EMBO Long-term Fellowship (ALTF 184-2019). S.W. is also supported by a Wellcome Principal Research Fellowship in Basic Biomedical Sciences (200846/Z/16/Z).

## STAR★Methods

### Key Resources Table

See separate file.

### Contact for Reagent and Resource Sharing

Further information and requests for resources and reagents should be directed to and will be fulfilled by the Lead Contact, Scott Waddell. Correspondence regarding the single-cell transcriptomic data and its analysis should be addressed to Vincent Croset.

### Experimental Model and Subject Details

#### Drosophila strains

GAL4 drivers used in this study are 0273-GAL4 (Burke et al., 2012; Gohl et al., 2011), R66C08-GAL4 (MBON-γ5β′2a) (Aso et al., 2014), repo-GAL4, tub-GAL80ts; repo-GAL4 (Lee and Jones, 2005; McGuire et al., 2003; Sepp and Auld, 1999), R86E01-GAL4 (Kremer et al., 2017) and R85G01-GAL4 (Kremer et al., 2017). UAS lines are w; +; UAS-mCherry, UAS-*aay*^RNAi^ (VDRC, 23179), UAS-*Inos*^RNAi^ (VDRC, 100763), UAS-*GstD1*^RNAi^ (VDRC, 103246), UAS-*CG33970*^RNAi^ (VDRC, 38661), UAS-*Socs36E*^RNAi^ (VDRC, 51821), UAS-*Irc*^RNAi^ (VDRC, 101098), UAS-*GstD9*^RNAi^ (VDRC, 103798), UAS-*Obp44a*^RNAi^ (VDRC, 43203), UAS-*Sod3*^RNAi^ (VDRC, 8760), UAS-*CG9377*^RNAi^ (VDRC, 42835), UAS-*Cyp28d1*^RNAi^ (VDRC, 7870), UAS-*Pcyt2*^RNAi^ (VDRC, 105794), UAS-*Drat*^RNAi^ (VDRC, 108325), UAS-*osy* (Wang et al., 2020), UAS-*GstD9* (FlyORF, F004171), UAS-*Obp44a* (FlyORF, F003929), UAS-*Sod3* (FlyORF, F003855), UAS-*aay* (FlyORF, F002296), UAS-*CG12338* (Dai et al., 2019), UAS-*Shi*^ts1^ (Kitamoto, 2001), UAS-*Kir2.1* (Paradis et al., 2001), UAS-*TNT* E (Sweeney et al., 1995), UAS-*GCaMP-7f* (Dana et al., 2019). Mutant strains are *NMDAR1*^K558Q^ and *NMDAR1*^F654A^ (Troutwine et al., 2019). Flies were raised on standard cornmeal food under a 12:12 light:dark cycle at 60% humidity and 25°C, unless otherwise stated. 3 day old mixed sex flies were used for single-cell transcriptomics, 3-6 day old mixed sex flies for water preference, RT-qPCR and imaging experiments, and 3-8 day old male flies for water consumption tests.

### Method Details

#### Water preference assays

Dehydration and water preference assays were performed as described (Lin et al., 2014b). Briefly, groups of 50-100 flies were stored for a given time period in vials containing a ∼2 cm layer of Drierite topped with a piece of cotton and a dried sucrose-coated filter paper. Vials were kept in a sealed container with a layer of Drierite at the bottom. Flies were (re-)hydrated flies in vials containing 1% agarose and a wet piece of sucrose-coated filter paper. After the appropriate water-deprivation period, flies were transferred into a T-maze and given the choice between two chambers lined with either a dried or a wet piece of filter paper. Preference Index was calculated as the number of flies in the wet tube minus the number of flies in the dry tube, divided by the total number of flies in each experiment.

#### Brain dissociation and cell collection

Brains (*0273-Gal4>mCherry*) were dissected and dissociated as described (Croset et al., 2018). Briefly, 24 central brains from an equal number of male and female flies were individually dissected in ice-cold DPBS (Gibco, 14190–086) and immediately transferred into 1 mL toxin-supplemented Schneider’s medium (tSM: Gibco, 21720–001 + 50 mM d(-)-2- amino-5-phosphonovaleric acid, 20 mM 6,7-dinitroquinoxaline-2,3-dione and 0.1 mM tetrodotoxin) on ice. Brains were washed once with 1 mL tSM and incubated in tSM containing 1.11 mg/mL papain (Sigma, P4762) and 1.11 mg/mL collagenase I (Sigma, C2674). Brains were washed once more with tSM and subsequently triturated with flame-rounded 200 mL pipette tips. Dissociated brains were resuspended in 1 mL PBS + 0.01% BSA and filtered through a 10 mm CellTrix strainer (Sysmex, 04-0042-2314). Cell concentration was measured using a disposable Fuchs-Rosenthal haemocytometer (VWR, 631–1096) under a Leica DMIL LED Fluo microscope. A typical preparation from 24 brains yielded ∼600,000 cells.

#### Library preparation, sequencing, and data processing

mRNA barcoding was performed using the Chromium Single Cell 3′ Reagent Kit v3 (10x Genomics), following the manufacturer’s instructions. For each sample, we targeted an 8,000 cell recovery. Two libraries were prepared for each condition (sated, 6h dehydrated, 12h dehydrated and 12h dehydrated + 45min rehydrated). Libraries were sequenced with NovaSeq 6000 (Illumina) at Oxford’s Wellcome Trust Centre for Human Genetics. We obtained 3.419 billion reads, and used CellRanger 3.1.0 to map these to the FB2018_06 *Drosophila melanogaster* genome assembly (v6.25) (Thurmond et al., 2019), and create digital gene expression (DGE) matrices.

#### Filtering and doublet removal

Cell barcodes with <300 or >4,500 features, >20,000 UMIs, >15% mitochondrial RNA, >10% rRNA or >15% ribosomal proteins were discarded. DGE matrices were then merged by condition, normalized, and scaled using SCTransform in Seurat v3 (Hafemeister and Satija, 2019; Stuart et al., 2019). This included regression for the effects replicate and sex – based on expression of the male-specific lncRNA *roX1* (Kelley and Kuroda, 2003). DGE matrices were integrated using CCA anchor-based methods, and cells were clustered using the Louvain algorithm on the shared nearest neighbor graph with a resolution of 2 and an UMAP reduction performed for visualization, using the top 20 Principal Components (PCs). Doublets were removed using a hybrid method. First, DoubletFinder (McGinnis et al., 2019) was run on data processed with the standard Seurat scaling method (without SCTransform), as we found that DoubletFinder failed to produce reliable BCmvn curves on SCtransformed data. This identified 3,493 potential doublets. Second, because the glial marker *nrv2* (Sun and Salvaterra, 1995) and markers for neurons releasing fast-acting neurotransmitters *VAChT*, *VGlut* or *Gad1* are normally not co-expressed, we considered it parsimonious to classify “cells” expressing two or more of these genes as doublets. So as not to needlessly discard cells containing small contamination from these genes, we set thresholds of 3 (*nrv2)*, 1.5 (*VAChT*), 2.1 (*VGlut*) and 2.3 (*Gad1),* above which we considered these genes to be highly expressed. These values were estimated from the normalized and scaled expression levels as the local minima in each gene’s bimodal distribution of non-zero values. We used the same principle to detect doublets in Kenyon Cells. We first identified Kenyon Cell clusters based on expression of *Dop1R1*, *ey* and *mub* (Croset et al., 2018). We then flagged all cells co-expressing markers for more than one Kenyon Cell subtype, namely *Ca-alpha1T* for *αβ*, *ab* for *γ* and *CG8641* for *α¢β¢* (Croset et al., 2018), with thresholds of 1.5, 1.5 and 2.2, respectively (again estimated as the local minima in each gene’s bimodal distribution of non-zero values in their normalized and scaled expression levels). Overall, this co-expression strategy identified another 2,828 doublets. Doublets were evenly spread across clusters, with only six clusters comprised of >20% doublets. All doublets identified with either of these two methods (9.05% of all cells) were removed prior to subsequent analyses (Figure S1A).

#### Clustering

We developed a 4-step clustering pipeline with the aim of minimizing the number of PCs used for clustering and reducing the ‘curse of high dimensionality’ effects on nearest neighbor search (Aggarwal et al., 2001; Bellman, 1961). After doublet removal, data was clustered using the Louvain algorithm with the first 20 PCs and a resolution of 2 and an UMAP plot constructed using the same PC dimensions for visualization. Then each cluster was assigned to one of six major cell types present in the *Drosophila* brain, using expression of the following markers: *VAChT* (cholinergic), *VGlut* (glutamatergic), *Gad1* (GABAergic), *ey*, *Dop1R2*, *Pka-C1*, *mub* (Kenyon Cells), *Vmat* (monoaminergic) and *CG10433* (glia & astrocytes) (Croset et al., 2018), with a seventh group containing non-assigned cells (“other”). Cells from each group were subsequently isolated. Data was normalized and scaled using SCTransform, and clustered and visualized on a UMAP plot, with 19, 17, 16, 12, 16, 14 and 12 PCs for each group, respectively. A resolution of 1 was used, except for the monoaminergic (4) and other (2) groups, with the aim of slightly over-clustering. Lastly, for each group a phylogenetic tree of all clusters was constructed in PCA space using Seurat’s BuildClusterTree function. Clusters on neighboring branches with less than 10 protein-coding genes differently expressed between them (Wilcoxon signed-rank test, adjusted p<0.05) were fused.

#### Annotation

Specific clusters were annotated based on the expression of the following known marker genes. Olfactory projections neurons (cholinergic): *acj6*, *ct*, *Lim1* (Komiyama and Luo, 2007; Komiyama et al., 2003; Lai et al., 2008), ellipsoid body (EB) large-field ring neurons (GABAergic): *cv-c*, *Dh31*, *Octbeta2R*, *5-HT7* (Becnel et al., 2011; Donlea et al., 2014; Kunst et al., 2014; Zhao et al., 2021), EB small-field ring neurons (GABAergic): *cv-c*, *Dh31*, *Octbeta2R*, not *5-HT7*, ventral and dorsal fan-shaped body (GABAergic): *cv-c*, *Dh31, sNPF*, not *Octbeta2R* (Nässel et al., 2008), medial fan-shaped body: *cv-c*, *Dh31*, not sNPF, not *Octbeta2R*, a/b Kenyon Cells: *sNPF*, *Eip93F* (Croset et al., 2018; Johard et al., 2008), g Kenyon Cells: *sNPF*, *ab* (Croset et al., 2018), a¢/b¢ Kenyon Cells: *CG8641* (Croset et al., 2018), dopaminergic neurons (monoaminergic): *ple*, *DAT* (Neckameyer and Quinn, 1989; Pörzgen et al., 2001), serotonergic neurons (monoaminergic): *SerT*, *Trh* (Coleman and Neckameyer, 2005; Demchyshyn et al., 1994), octopaminergic neurons (monoaminergic): *Tdc2*, *Tbh* (Cole et al., 2005; Monastirioti et al., 1996), tyraminergic neurons (monoaminergic): *Tdc2*, not *Tbh*, astrocytes: *AANAT1*, *alrm*, *Gat*, *e* (Davla et al., 2020; Doherty et al., 2009; Muthukumar et al., 2014), surface glia: *Tret1-1*, *Mdr65* (DeSalvo et al., 2014; Volkenhoff et al., 2015), ensheathing glia: *zyd*, *trol* (Konstantinides et al., 2018; Melom and Littleton, 2013), cortex glia: *zyd*, *wrapper* (Konstantinides et al., 2018). Thresholds for each gene were set manually. Top markers for each cluster were calculated (Data S1).

#### Differential expression (DE)

To account for zero-inflation triggered by dropout events and enable the use of DE tools initially created for analyzing bulk RNA-sequencing data, we generated cell-specific weights, using ZINB-WaVE (Risso et al., 2018; Van den Berge et al., 2018). Both DESeq2 (Love et al., 2014) and edgeR (McCarthy et al., 2012; Robinson et al., 2010) were then used to calculate differential expression between pairs of conditions. Genes with |log2(FC)|>1 and adjusted p-value<0.05 calculated with either method were considered to be differentially expressed.

#### Pathway analysis

Gene Ontology Biological process over-representation test was performed for the differentially expressed genes shown in Figure 2E, using the Bioconductor package ClusterProfiler (Yu et al., 2012). Significantly enriched GO terms based on p-values were then visualized as a chord diagram with GOplot (Walter et al., 2015).

#### RT-qPCR

Quantitative PCR experiments were performed as described (Croset et al., 2018). In brief, total RNA was extracted from groups of 40 fly heads using the RNeasy Mini kit (Qiagen 74104). mRNA was then reverse-transcribed using the SuperScript III First-Strand Synthesis SuperMix (Invitrogen, 18080400) according to manufacturer’s instructions. qPCR was performed in a Light-Cycler 480 Instrument II (Roche, 05015243001) using the Universal Probe Library system (UPL; Roche, 04683633001 and 04869877001). Each 10 mL reaction contained 2.4 mL of pre-amplified cDNA, 0.4 mM of each primer (designed with Roche Assay Design Center), 0.2 mM of UPL probe, and 5 mL LightCycler 480 Probes Master (Roche, 4887301001). Cycles were as follows: 95 °C, 10’; 45X [95 °C, 10’; 60 °C, 30’; Fluorescence acquisition; 72 °C, 1’].

#### D-serine Feeding

For D-serine feeding 2.9 g/L or 27.6 mM of D-serine (S0033 Tokyo Chemical Industry) was mixed with 2% agarose and 5% sucrose. We used the same concentration of L-serine (S0035 Tokyo Chemical Industry). For control treated groups we used 2% agar and 5% sucrose.

#### CAFE Assay

CAFE assay was performed as described (Park et al., 2018) but adjusted for water consumption. Groups of 12 male flies were first dehydrated for 1 h in vials containing Drierite without indicator (ACROS Organics). Flies were then transferred to CAFE vials containing four glass capillaries loaded with 5 µL of deionized water and ∼0.1 µL of mineral oil. Vials were loaded into a tray covered with plastic wrap to reduce water loss due to evaporation. We also used 3 empty CAFE vials to assess the level of evaporation to subtract from the amount of water consumed by the flies. Flies were kept in either 20°C or 30°C incubators. Food consumption CAFE used 5% yeast extract and 5% sucrose solution and 10 male flies per vial measuring consumption across 24 h. Vials also contained 10 mL of 1% agarose. We kept track of the number of deaths for each vial and re-adjusted consumption values based on the number of flies alive by the conclusion of the assay. Water and food consumption in µL is recorded as total water consumption of 12 flies minus the average evaporation volume from the empty vials.

D-serine preference CAFE involved storing groups of 8 male flies per vial at 30°C for 18-20 h prior to the start of the assay. We then transferred the flies into CAFE preference vials containing two capillary tubes loaded with normal 5% yeast extract and 5% sucrose and two capillary tubes loaded with 2.9 g/L (27.6 mM) of D-serine (S0033 Tokyo Chemical Industry) with 5% yeast extract and 5% sucrose. Vials also contained 10 mL of 1% agarose. Flies were then given 24 h to feed then were removed. D-serine Preference Index was calculated as the volume of D-serine food minus volume of normal food divided by the total volume of food consumed.

#### Manual Feeding Assay

The water feeding assay was performed as described (Jourjine et al., 2016) with minor adjustments. Flies were briefly anesthetized on ice and fixed to a glass slide using beeswax. The slide was then transferred to an empty pipette tip box containing Drierite without indicator (ACROS Organics), closed, then wrapped with parafilm at the seam. For experiments performed at 30°C we put the flies in an incubator for the 2 h dehydration step then performed the experiment in a temperature (30°C) and humidity-controlled (60-65%) booth. The flies were hand fed with a syringe containing deionized water. If the experiments were performed at 30°C the water was warmed to minimize temperature changes in the fly during feeding. We presented the water 10 times to each fly and measured cumulative time spent drinking the water. Presentations were made such that we briefly touched the fly’s legs with the water. Flies were excluded if they exhibited little to no movement when touched on their abdomen with the needle. For NMDAR single-site mutant flies we used a 2.5 h dehydration step instead of 2 h as the background Canton-S strain from the Atkinson lab (UT, Austin) showed greater dehydration resistance than Canton-S strain of the Waddell lab.

#### Hygrosensation/Water memory in the T-maze

We used 3-5 day old flies to assess hygrosensation in the water T-maze, as described (Sayeed and Benzer, 1996). Briefly, 24 h prior to testing we group housed ∼100 flies per vial on standard cornmeal/agar food and a 20x60 mm piece of filter paper. If flies were water deprived, they were housed in vials containing desiccant for 2 h before testing. Testing was performed in a temperature (30°C) and humidity-controlled (60-65%) booth. Flies were loaded into a T-maze and given a 2 min choice between an arm containing wet filter paper or dry filter paper. Preference Index was calculated as the number of flies in the wet arm minus the number in the dry arm, divided by the total number of flies.

For water memory assays we used 3-5 day old flies. Prior to training, flies were water deprived for 16-17 h in vials containing dessicant. Water-deprived flies were trained to associate odor with water reward (Lin et al., 2014a) then were immediately transferred to vials containing 1% agarose (as a water source) supplemented with nothing, D-serine, or L-serine, 29 g/L (276mM). They were then housed for an additional 23 h in vials containing dry sucrose, dry sucrose and D-serine, or dry sucrose and L-serine, before being tested in the T-maze for water memory performance.

#### Two-Photon Calcium Imaging

Imaging experiments were performed using 3-7 day old flies as described previously (Jacob and Waddell, 2020; Owald et al., 2015). Briefly, flies were immobilized by cooling on ice and mounted in a custom-built chamber to allow free movement of their legs. For control conditions we used an external saline containing 103 mM NaCl, 3 mM KCl, 5 mM N-Tris, 10 mM trehalose, 10 mM glucose, 7 mM sucrose, 26 mM NaHCO_3_, 1 mM NaH_2_PO_4_, 1.5 mM CaCl_2_, 4 mM MgCl_2_, osmolarity 275 mOsm, pH 7.3. The head capsule was opened under room temperature carbogenated (95% O_2_, 5% CO_2_) external saline. The mounted fly was placed under the two-photon microscope (Scientifica). We used a Ti-Sapphire laser (Chameleon Ultra II, Coherent) to excite fluorescence using 140 fs pulses with 80 MHz repetition rate at 910 nm. We acquired 256 x 256 pixel images at 5.92 Hz controlled by ScanImage 3.8 software (Pologruto et al., 2003).

For analysis, the two-photon images were manually segmented using Fiji (Schindelin et al., 2012). With a custom Fiji script we measured the average baseline fluorescence (F_0_) for 14 s prior to each drug treatment or stimulus delivery. F/F_0_ describes the fluorescence relative to the baseline. For drug delivery treatments we defined the “Pre-” treatment as the average F/F_0_ value for 14 s prior to the drug delivery and the “Post-” treatment as the average F/F_0_ in the 25 s from onset of drug delivery to the offset. To account for inter-cell baseline differences in F/F_0_ we normalized the “Pre-” treatment to equal 0 for each cell. For area under the curve (AUC) calculation we measured the approximated integral of F/F_0_ during the drug treatment (“During”) as well as after the treatment until the recording ended (“After”). To account for variance between individual cells, we normalized the beginning of the trace starting from the onset of drug delivery to equal 0 for each cell. For odor delivery AUC analysis we measured the F/F_0_ during the 5s odor presentation.

To image fly brains following desiccation or starvation, we starved and water deprived flies for 8-10 h prior to recordings. All recordings were performed between ZT 2-6 and deprived flies at ZT 18-22. Dehydration was performed as above.

#### Solutions

For acute drug application, we used a perfusion pump system (Fisher Scientific US 14-284-201) to continuously deliver saline at a rate of ∼0.043 mL/sec. All drugs were applied with in the presence of 1µM tetrodotoxin (TTX) to block voltage-gated sodium channels and propagation of action potentials.

#### NR1+ neuron and astrocyte recordings

Recordings of NR1+ neurons (*nmdar1*-*KIGAL4*>*GCaMP7f*) were performed using either 4 mM Mg^2+^ or 0 mM Mg^2+^ saline. To compensate for the change in osmolarity with Mg^2+^ free saline, we added 1.5 mM CaCl_2_ and 2.5 mM NaCl. We used 20 mM N-methyl-D-aspartic acid (Sigma-Aldrich M3262), D-serine (Tokyo Chemical Industry - S0033), L-serine (Tokyo Chemical Industry – S0035), and glycine (Sigma-Adlrich G8898). Drug mixtures were maintained at room temperature prior to application. Before recordings we pre-loaded the pump with 1 mL of saline then 1 mL of drug mixture and allowed the pump to run continuously throughout the recording so there were no discontinuities in flow during recording. We flushed the pump for 5 min with saline between each drug application. When multiple drug treatments were performed the application order was randomized for each animal to account for sensitization/desensitization effects. To block NMDARs, we applied 15 mM ketamine with NMDA, D-serine, and TTX.

Astrocyte recordings (*R86E01-GAL4*>*GCaMP7f*) were performed using normal external saline, hyperosmotic 320 mOsm saline (Jourjine et al., 2016), and sugar free saline (Cheriyamkunnel et al., 2021). Hyperosmotic saline was made by adjusting the total amount of added solute in our original saline recipe, while maintaining the same ratios, so that the osmolarity was increased to 320 mOsm. We applied 10 mM acetylcholine (Sigma-Aldrich A6625), 10 mM ATP (Sigma-Aldrich A2383), 10 mM glutamate (Sigma-Aldrich G0355000), or 10 mM adenosine (Sigma-Aldrich A9251) with 1 µM TTX (Sigma-Aldrich T3194). When multiple drug treatments were performed the application order was randomized for each animal.

For MBON-γ5β′2a (M6) recordings (*R66C08-GAL4*>*GCaMP7f*), flies were exposed to a constant air stream containing mineral oil solvent (air). Flies were sequentially exposed to 3-octanol (OCT) and 4-methylcyclohexanol (MCH) for 5 s with 30 s inter-odor interval. To measure dendritic responses of MBON-γβ′2a signals were simultaneously acquired from both hemispheres and average responses were analyzed.

For tetanus toxin (TetX) application, we replaced the bath saline with saline containing 1 µM TetX and incubated the flies for 1 h at room temperature. Halfway through the incubation the bath solution was mixed 4X by pipetting.

For α,β-methylene adenosine 5’-diphosphate (AMPCP) (Sigma-Aldrich M3763) application we incubated the flies with 100 µM AMPCP in normal saline for at least 30 min. Flies were then imaged using saline containing AMPCP and ATP.

#### Identification of astrocytes and review of EM data reconstructions

We used FlyWire (www.flywire.ai) (Dorkenwald et al., 2022) to construct the morphology of astrocytes in the Full Adult Female Brain (FAFB) transmission electron microscope (TEM) dataset (Zheng et al. 2018). Astrocytes were identified and distinguished from other glia in the raw EM data by first following cell processes that cross the ensheathing glial boundary, and following them to find their cell body. Astrocytic morphology is characteristically distinct from that of neuronal soma tracts, and ensheathing or cortex glia.

In FlyWire, 3D meshes from automatic A.I. based (Lee et al., 2017) reconstructions of neurons and glia are readily available, but manual expert review is required to add missing, or remove extraneous, branches. Following revision neurons and glial cells had no false continuations, by standard criteria that are previously described (Dorkenwald et al., 2022; Otto et al., 2020; Schneider-Mizell et al., 2016). In brief, an expert reviewer (>1000 h experience) first inspected the astrocytes for false continuations, and obvious missing branches. Then a methodical visual inspection of all branches with neuroglancer (Dorkenwald et al., 2022) allowed the user to identify and correct further smaller errors. A second expert reviewer then performed the same review to generate a consensus. In this study, 4 astrocytes were submitted for extensive review to two experienced reviewers following every process section by section from the tip inwards towards. This review process only resulted in the correction of a few smaller branches with no more than 2 connections each.

#### Classification of astrocytic processes into tripartite synapses vs. bouton engulfing

Tripartite synapses (TPS) contain presynaptic, postsynaptic, and astrocytic process, with all compartments making direct contact with the synaptic cleft. We found that astrocytic processes sometimes also cover a large part of the perimeter of a presynaptic bouton without contacting the synaptic cleft – a motif we classified as ‘engulfing’. Lastly astrocytic processes were often also found to contact the post-synaptic neuron below the synaptic cleft (within 500 nm of a presynaptic terminal. These contacts to post-synaptic neurons were not included in the TPS analyses.

#### Analysis of astrocyte-neuronal type relationship

3D meshes of cells in FlyWire, information about synapses with pre and postsynaptic partners (Buhmann et al., 2021), and neurotransmitter predictions (Eckstein et al., 2020) were downloaded with the CAVEclient (github.com/seung-lab/CAVEclient) and further analysed using functionalities from FAFBseg v.1.4 (https://github.com/navis-org/fafbseg-py) and Navis v.0.6 (https://github.com/navis-org/navis) and custom scripts (available upon request). Synapses with CleftScore ≥ 50 and ConnectionScore ≥ 33 (Buhmann et al., 2021; Heinrich et al., 2018) were considered for analyses, after thresholds were manually reviewed in a sample of 500. Synapses were classified as TPS, when an astrocyte was found as a postsynaptic partner to a neuronal presynapse.

To determine the vicinity profile of an astrocyte, it’s mesh was obtained and skeletonised with trimesh v.3.9.32 (https://github.com/mikedh/trimesh) and Skeletor 1.1 (Au et al., 2008).

To exclude somata and main branches we only considered nodes with a radius of < 300 nm. To generate the vicinity profile, synapses within a 2 µm bounding box around the nodes were filtered for the above criteria and classified by neurotransmitter usage. Then the histogram of distances between the nodes and synapses of each neurotransmitter was generated and gaussian kernel density estimates and bootstrap samples of the means of the distance distributions computed with Python base functions for statistical analyses (Code available on request).

#### 3D representations

3D representations and animations of astrocytes were created with Blender 3.0 (Blender Community, 2018) from meshes obtained from Flywire, via CAVEclient and neuroglancer within Flywire.

### Code availability

Detailed code for scSeq analyses is available at https://github.com/sims-lab/FlyThirst.

## Supplementary Figure Legends

**Figure S1. Filtering and clustering single-cell transcriptomics data, Related to Figure 1.**

(**A**) Left: UMAP plot obtained after initial clustering, showing doublets identified with DoubletFinder or the co-expression strategy. Right: barplot showing the number of cells labelled with either method and that were discarded prior to subsequent analyses. (**B**) Heatmap showing expression of the top marker genes from six of the main cell classes. (**C**) Number of cells per cluster, grouped by main cell classes. (**D**) Same UMAP plot as Figure 1E, with colors representing the experimental sample from which each cell originates. (**E**) Ratio of cells originating from each sample, for each cluster, ordered by size. All clusters contain cells from each condition and sample. As expected, more variability was evident in the small clusters. (**F**) Heatmap showing expression of the top marker genes from each glia cluster.

**Figure S2. Differential gene expression in thirsty flies, Related to Figure 2.**

(**A**) Venn diagrams showing the number of gene expression events identified with DESeq2, edgeR, or both methods. Top: up-regulated genes. Bottom: down-regulated genes. (**B**) Relative expression changes for a selection of differentially expressed genes in thirsty flies. Groups are those as in Figure 1B, and values are relative to those in sated flies. Data are from RT-qPCR of whole heads, n2-3. (**C**) Relative expression changes for a selection of genes that were differentially expressed in thirsty flies, in a 12h dehydrated and 12 h starved condition, relative to levels in sated animals. RT-qPCR on whole heads, n2-3. (**D**) Chord diagram showing the most represented GO categories across the genes that are differently expressed in glia of thirsty flies.

**Figure S3. Additional water consumption and water preference data, Related to Figure 3.**

(**A**) CAFE water consumption assay protocol. Orange section of black line indicates period when flies are water deprived. (**B**) Knockdown of *Drat* in glia does not change water consumption in CAFE. (**C**) Knockdown of *inos* in glia does not alter water consumption in CAFE. (**D**) Knockdown of *aay* in glia reduces water consumption in CAFE. (**E**) Overexpression of *aay* in glia increases water consumption in CAFE. (**F**) Knockdown of *aay* does not alter food consumption as measured in CAFE. (**G**) Number of deaths per vial of 12 flies from S1D during CAFE assay shows overexpression of *aay* makes flies more susceptible to water deprivation. (**H**) Flies with *aay-*knockdown in glia exhibit normal hygrosensation in the T-maze. Flies show normal innate avoidance of high humidity when water sated, and display normal water seeking when water deprived. (**I**) Temperature regimen for inhibiting astrocyte vesicle release with expression temperature sensitive *Shibire^ts^* while measuring water consumption. (**J**) S*hibire^ts^* expression in astrocytes does not significantly suppress water consumption, relative to controls. (**K**) Overexpressing the inward-rectifying potassium channel Kir2.1 in astrocytes did not significantly change water consumption. (**L**) Permissive temperature control for *R86E01>TrpA1* activation experiments (Figure show that flies that do not have activated astrocytes do not increase water consumption (n = 14, 15 flies). (**M**) UMAP plot showing expression levels of *Syb* across glial clusters. Scale represents normalized expression levels. Refer to Figure 1F for annotations. (**N**) UMAP plot showing expression levels of *Syx1A* across glial clusters. Scale represents normalized expression levels. Refer to Figure 1F for annotations. For panels B-H and J-K, normality was assessed using the Shapiro-Wilk test. * p<0.05, ** p<0.01, Ordinary one-way ANOVA with Dunnett’s multiple comparisons test, unless otherwise stated. Data are mean +/- SEM.

**Figure S4. Effects of D-serine feeding and D-serine preference, Related to Figure 4.**

(**A**) Protocol for D-serine feeing and testing in the CAFE assay and results from 1 and 3 days of D-serine feeding. Only 3 days of D-serine feeding significantly increased water consumption. * p<0.05 two-tailed Mann-Whitney test (n_1-day_ = 10 vials, n_3-days_ = 16 vials). (**B**) Neither *aay*-knockdown or GAL4/GAL80^ts^ control flies show a preference for D-serine containing food in CAFE (n = 13, 14 vials). (**C**) D-serine feeding does not change food consumption in the CAFE (n = 9 vials). (**D**) Survival of *aay*-knockdown flies in CAFE at permissive 20°C (no induction of *aay*-RNAi expression) that were either fed D-serine or kept on ‘D-serine free’ food. Flies survived throughout the assay (n = 8, 9 vials). (**E**) Survival of *aay*-knockdown flies in CAFE with a 30°C heat step (induced *aay*-RNAi expression). Flies with *aay* knockdown that were also fed D-serine show significantly compromised survival in CAFÉ (n = 8, 9 vials). ** p<0.01 *aay*-RNAi vs. *aay*-RNAi + D-serine. Normality was assessed using the Shapiro-Wilk test. * p<0.05, ** p<0.01, *** p<0.001, **** p<0.0001 Ordinary one-way ANOVA with Dunnett’s multiple comparisons test, unless otherwise stated. Data are mean +/- SEM.

**Figure S5. D-serine is a co-agonist for NMDARs, Related to Figure 5**

(**A**) UMAP plot showing expression levels of *Nmdar1* across glial clusters. Scale represents normalized expression levels. Refer to Figure 1E for annotations. (**B**) UMAP plot showing expression levels of *Nmdar2* across glia clusters. Scale represents normalized expression levels. Refer to Figure 1E for annotations. (**C**) Individual ΔF/F_0_ GCaMP7f traces of NR1- expressing neurons to D-serine and NMDA application in high Mg^2+^. (**D**) Individual traces of NR1 neurons to glycine and NMDA application in high Mg^2+^. (**E**) Individual traces of NR1 neurons to D-serine and NMDA application in 0 mM Mg^2+^. (**F**) Individual traces of NR1 neurons to glycine and NMDA application in 0 mM Mg^2+^. (**G**) Individual traces of NR1 neurons to L-serine and NMDA application in 0 mM Mg^2+^. (**H**) Individual traces of NR1 neuron responses to D-serine and NMDA application, with and without ketamine. (**I**) quantification of Pre- and post-responses of Figure 5H with outliers removed. ** p<0.01, Kruskal-Wallis ANOVA with Dunn’s multiple comparisons test. (**J**) 3D representation of 11 reviewed astrocytes with processes in the SMP. Brain neuropil is grey background. Astrocyte somata reside at the neuropil boundary and their processes are mostly mutually exclusive to one area of the neuropil. Few astrocyte processes venture past ensheathing glial boundaries. Inset shows lesser zoom for position of reference within the entire brain. (**K**) Astrocyte cell bodies are often close to trachea. Left: Greyscale EM data with a glial soma (blue) and two tracheal branches (orange). Upper right: Same greyscale image with 3D reconstructions of astrocyte cell body and trachea superimposed. Lower right: 3D renderings show the astrocyte in relation to trachea. (**L**) Same EM section as Figure 5I with superimposed 3D reconstruction of neuronal boutons and astrocytic processes engulfing a bouton (top left) and contributing to a tripartite synapse (TPS) (top right). Bottom left: 3D renderings show astrocytic processes engulf the synaptic bouton near the synapse (asterisk). Bottom right: 3D reconstructions show 3 boutons are contacted by thin astrocytic processes. Location of synapse from Fig 5I indicated (asterisk). Scale bars 300nm. See also Video S1. (**M**) Astrocytic processes are closer to glutamatergic synapses, than GABAergic and cholinergic synapses. This is shown in a vicinity profile for SMP astrocytes consisting of histograms of the distribution of distances between synapses and fine astrocytic processes (less than 300nm radius), sorted by predicted neurotransmitter type. Note: Engulfment happens typically within 500nm of synapses and astrocytess often also contact postsynaptic processes more distant from the synapse. (**N**) Astrocytes are significantly closer to glutamatergic than cholinergic synapses. Means of 10000 bootstrap samples taken from the probability distributions of distances between synapses and glial processes of the SMP astrocytes. Only glutamatergic and cholinergic synapses in the vicinity of fine astrocytic processes (2µm radius around processes with radius *≤* 300nm) are displayed together with random draws from both sets. All 3 distributions are significantly different from each other. Heteroscedastic t-test: p = 0 (Glut <> Random) and p = 10^-200^ (ACh <> Random).

**Figure S6. Specificity of astrocytic responses to neurotransmitters, Related to Figure 6**

(**A**) Astrocytes show matched responses to ATP and ACh application. ΔF/F_0_ responses for ACh plotted against ATP responses with linear regression. Pie chart shows number of astrocytes with matched and mismatched responses between ACh and ATP. Fisher’s exact test matched vs. mismatched, p = 1.076 *10^-5^ OR = 3.28. (**B**) Astrocytes show matched responses to glutamate and ACh application. ΔF/F_0_ responses for glutamate plotted against ACh responses with linear regression. Pie chart shows number of astrocytes with matched and mismatched responses between glutamate and ACh. Fisher’s exact test matched vs. mismatched, p = 2.27 *10^-6^ OR = 3.72. (**C**) Schematic of two possible mechanisms for differential astrocytic responses with experimental procedure. 1) Exogenously applied neurotransmitter (red dots) acts on pre-synaptic or any non-astrocytic compartment that drives release of a secondary neurotransmitter (green dots, which then drives changes in astrocytic Ca^2+^. 2) Exogenously applied neurotransmitter (red dots) acts directly on the astrocytes modulating Ca^2+^ levels. Co-applying tetanus toxin with exogenous neurotransmitter application should prevent mechanism 1) from occurring. (**D**) 1 h of 1µM Tetanus toxin (TetX) application is sufficient to suppress polysynaptic activity. Odor exposure (black bar) to the fly activates MBON-γ5β′2a (M6). Tetx suppresses odor-induced activity recorded in the β′2a region of the MBON-γ5β′2a dendrite. (**E**) Area Under the Curve (AUC) analysis of β′2a dendritic field before and after Tetx application shows elimination of odor-induced Ca^2+^ response. ** p<0.01 paired t-test. (**F**) Most glutamate and ATP responsive astrocytes retain their neurotransmitter sensitivity following Tetx application. Tetx application reduced the number of ATP sensitive astrocytes and modestly reduced glutamate responsive astrocytes (p = 0.066). * p<0.05 Fisher’s exact test with Bonferroni correction. (**G**) Protocol for ATP and adenosine application, and ATP with AMPCP. (**H**) Astrocytes show excitatory responses to ATP but not to adenosine. ** p<0.01, Kruskal-Wallis ANOVA with Dunn’s multiple comparisons test. (**I**) Astrocytes show inhibitory responses to ATP and adenosine. * p<0.05, ** p<0.01, Kruskal-Wallis ANOVA with Dunn’s multiple comparisons test. (**J**) Inhibiting ATP degradation by blocking ecto-5′-nucleotidase/CD73 with AMPCP does not abrogate ATP responses ** p<0.01, Kruskal-Wallis ANOVA with Dunn’s multiple comparisons test.

**Figure S7. Astrocytic responses to ATP and glutamate in hungry and thirsty flies, Related to Figure 7**

(**A and B**) Astrocytes show differential responses to ATP and glutamate in hungry flies. Bold lines are mean responses and shade is SEM. Pie charts show proportion of astrocytes that show positive, no change (NC), and negative responses to ATP and glutamate. (**C**) Quantification of ΔF/F_0_ before and after neurotransmitter application in hungry flies. ** p<0.01, Kruskal-Wallis ANOVA with Dunn’s multiple comparisons test. (**D**) Ca^2+^ imaging traces of positive responsive astrocytes in thirsty and sated animals with ATP application. Line is mean and fill SEM. (**E**) Violin plot showing the difference of *aay* expression in all astrocytes in sated and 12h dehydrated flies. Each dot represents an individual cell. (F) Area Under the Curve (AUC) of excitatory ATP responses of astrocytes in sated vs. thirsty animals. ** p<0.01, Kruskal-Wallis ANOVA with Dunn’s multiple comparisons test. (G) Water deprivation desynchronizes matched astrocyte responses for glutamate and ATP. ** p<0.01, Fisher’s exact test with Bonferroni correction.

**Video S1. Fine astrocytic processes often engulf glutamatergic synaptic boutons or contribute to tripartite synapses, Related to Figures 5I and S5L.** 3D representations of an astrocyte (blue) in the SMP and glutamatergic neurons (green) with boutons containing annotated presynapses (red), which consist of synaptic vesicles (red spheres) and pre-synaptic T-bar densities (black). 00:00 to 00:55, When glial cells engulf axons, multiple processes of different sizes contact a substantial area of a presynaptic bouton’s membrane but do not extend into the synapse. 00:56-1:40, Fine astrocyte processes either pass through the synaptic cleft, or terminate within it, to form a tripartite synapse. In both cases the astrocyte processes contact the synaptic cleft alongside the postsynaptic neurons. Many closely spaced boutons of the same presynaptic neuron can be contacted by the same astrocyte, which therefore contributes processes to multiple tripartite synapses. Anatomical features are further highlighted in the video. Cellular meshes were retrieved from FlyWire and animations produced with blender.

